# Giant genes are rare but implicated in cell wall degradation by predatory bacteria

**DOI:** 10.1101/2023.11.21.568195

**Authors:** Jacob West-Roberts, Luis Valentin-Alvarado, Susan Mullen, Rohan Sachdeva, Justin Smith, Laura A. Hug, Daniel S. Gregoire, Wentso Liu, Tzu-Yu Lin, Gabriel Husain, Yuki Amano, Lynn Ly, Jillian F. Banfield

## Abstract

Across the tree of life, gene lengths vary, but most are no more than a few thousand base pairs in length. The largest protein often reported is the ∼40,000 aa eukaryotic Titin. Even larger proteins may occur in the rapidly expanding set of metagenome-derived sequences, but their existence may be obscured by assembly fragmentation. Here, we leverage genome curation to complete metagenome-derived sequences that encode predicted proteins of up to 85,804 aa. Overall, the findings illuminate a huge knowledge gap related to giant proteins. Although predicted proteins of >30,000 aa occur in bacterial phyla such as *Firmicutes* and *Actinobacteria*, they are most common in ca. *Omnitrophota,* ultra small bacteria that adopt predatory lifestyles. All full length giant genes encode numerous transmembrane regions and most encode divergent secA DEAD helicase domains. *In silico* structural prediction of protein subregions was required to identify domains in unannotated protein segments, and revealed putative domains implicated in attachment and carbohydrate degradation. Many giant genes in new complete and near-complete *Omnitrophota* genomes occur in close proximity to genes homologous to type II secretion systems as well as carbohydrate import systems. This, in combination with the domain content, suggests that many bacterial giant proteins enable prey adhesion and cell wall digestion during bacterial predation.

## Introduction

It has been ∼15 years since the last evaluation of the incidence of large bacterial genes (≥ 25 kb) and their protein products <u>(Reva and Tümmler 2008)</u>. Recently it was reported that the genomes of some ca. *Omnitrophica* (formerly also referred to as WOR-2, OP3, but henceforth *Omnitrophota*) encode open reading frames (ORFs) of ≥ 20 kbp in length (≥ 7,000 amino acids) that are transcribed on a single mRNA (Seymour et al. 2023). The authors observed a 3-fold increase in the abundance of a 39,678 amino acid protein when *V. archaeovorus LiM* was attached to its prey organism, *Methanosaeta,* via proteomics. This, in combination with glycosyl hydrolase, glycosyl transferase, hydrolase, and peptidase domains, suggests a role for giant *Omnitrophota* proteins in predation (Kizina et al 2022). This is consistent with prior observations suggesting a predatory lifestyle for *Omnitrophota* <u>(Perez-Molphe-Montoya et al. 2022) (Seymour et al. 2023)</u>. One representative, *ca. Velaminicoccus archaeovorus sp. LiM,* predates specifically on *Methanosaeta,* archaea of the family *Methanosarcinales*, in co-culture <u>(Kizina et al. 2022)</u>. These findings motivated a targeted search through metagenomic data to find additional giant genes, both within *Omnitrophota* and in genomes of other Bacteria and Archaea.

Predation by microorganisms is a major force influencing the structural dynamics of microbial communities, along with nutrient cycling and energy transfer. Predation has been identified as the lifestyle of various isolated bacteria belonging to the phyla *Bdellovibrionota* <u>(Van Essche et al. 2011)</u>, *Myxococcota* <u>(Thiery and Kaimer 2020)</u>, *Planctomycetota* <u>(Shiratori et al. 2019)</u>*, Actinobacteriota* <u>(Kumbhar et al. 2014)</u>, *Chloroflexota* <u>(Livingstone et al. 2018)</u>, and *Proteobacteria* <u>(Rashidan and Bird 2001) (Gupta et al. 2016)</u>. Notable representatives include *Bdellovibrio bacteriovorus* (*Bdellovibrionota*), and *Micavibrio aeruginosavorus* (*Alphaproteobacteria*). Observed modes of predation between these two groups of predatory bacteria differ in that *Bdellovibrio* reproduces inside the host cell prior to host lysis whereas *Micavibrio* displays an epibiotic mode of predation <u>(Davidov et al. 2006)</u>. Evidence of predation by nanobacterial *Absconditabacteria*, a lineage of the Candidate Phyla Radiation (CPR), has been documented in recent studies <u>(Moreira et al. 2021) (Bor et al. 2018)</u>. In this case, the prey are other bacteria. These findings raise the question of the mechanisms by which predatory bacteria kill and digest their prey.

Here, we leverage public and newly generated datasets, including newly curated complete genome sequences, to reexamine the size range and phylogenetic distribution of very large genes and to test the hypothesis that giant proteins are widespread and potentially a general mechanism for predation. We categorize giant proteins and investigate their domain content and thus possible functions and potential origins. Protein functional analyses relied upon detailed bioinformatics-based *in silico* structural predictions because biochemical studies of giant proteins are exceedingly difficult and very low throughput. Our analyses bring to light a remarkable incidence of large proteins in *Omnitrophota.* Domain analyses strongly suggest that the protein product or products strongly associate with the Omnitrophota cytoplasmic membrane and enable attachment and degradation of prey cell walls to generate substrates that are used in glycolysis. The ubiquitous presence within giant proteins of protease domains involved in autocatalytic cleavage of precursor peptide sequences suggests post-translational cleavage of these proteins into multiple protein products. *In silico* protein structure prediction of these giant proteins results in several large regions with regular, periodic folding structures that show structural homology to carbohydrate binding modules implicated in cell wall adhesion, as well as novel long beta helix domains. Overall, we find that large genes, although rare, are near-ubiquitous over a restricted phylogenetic distribution, can achieve remarkably large sizes, and are likely key to predatory lifestyles of *Omnitrophota*, in which they occur with the greatest average frequency.

## Results and Discussion

### Size and distribution of giant proteins in bacteria and archaea

We leveraged vast public databases of genome sequences, most of which derive from metagenomes and thus capture sequences from uncultivated microorganisms, to evaluate the size range and taxonomic distribution of very large genes. We refer to genes of > 30,000 bp in length (10,000 aa) as encoding “large” proteins, and genes of > 90,000 bp (≥ 30,000 aa) as encoding “giant” proteins.

Compared to genomic representatives for other phyla from the GTDB database, we find that *Omnitrophota* genomes contain the highest average copy number of very large genes per genome (**Fig. 1**). Very large genes encoding proteins ≥ 10,000 aa are frequently detected in other phyla within the PVC supergroup such as *ca. Elusimicrobiota, ca. Hinthialibacteriota, Verrucomicrobiota*, *ca. CG03* and *Planctomycetota*. Notably, predicted proteins of length ≥ 5,000 aa are absent from all 34 *Ratteibacteria* genomes in this dataset. Other non-PVC phyla with high average incidence of very large genes are *Actinobacteriota, Myxococcota*, and *Margulisbacteria* (**Fig. 1b**). A caveat is that many of the genomes in public databases are fragmented, reducing the overall incidence of very large ORFs. For Archaea we found many fewer genes encoding proteins > 10,000 amino acids relative to bacteria, with the largest number of such sequences observed in the Asgard supergroup (**Figure 1c**). Notably, sequences encoding proteins of >30,000 aa are not observed in any other phylum within the PVC supergroup other than *Omnitrophota* (**Fig. 1a**).

**Fig. 1:**
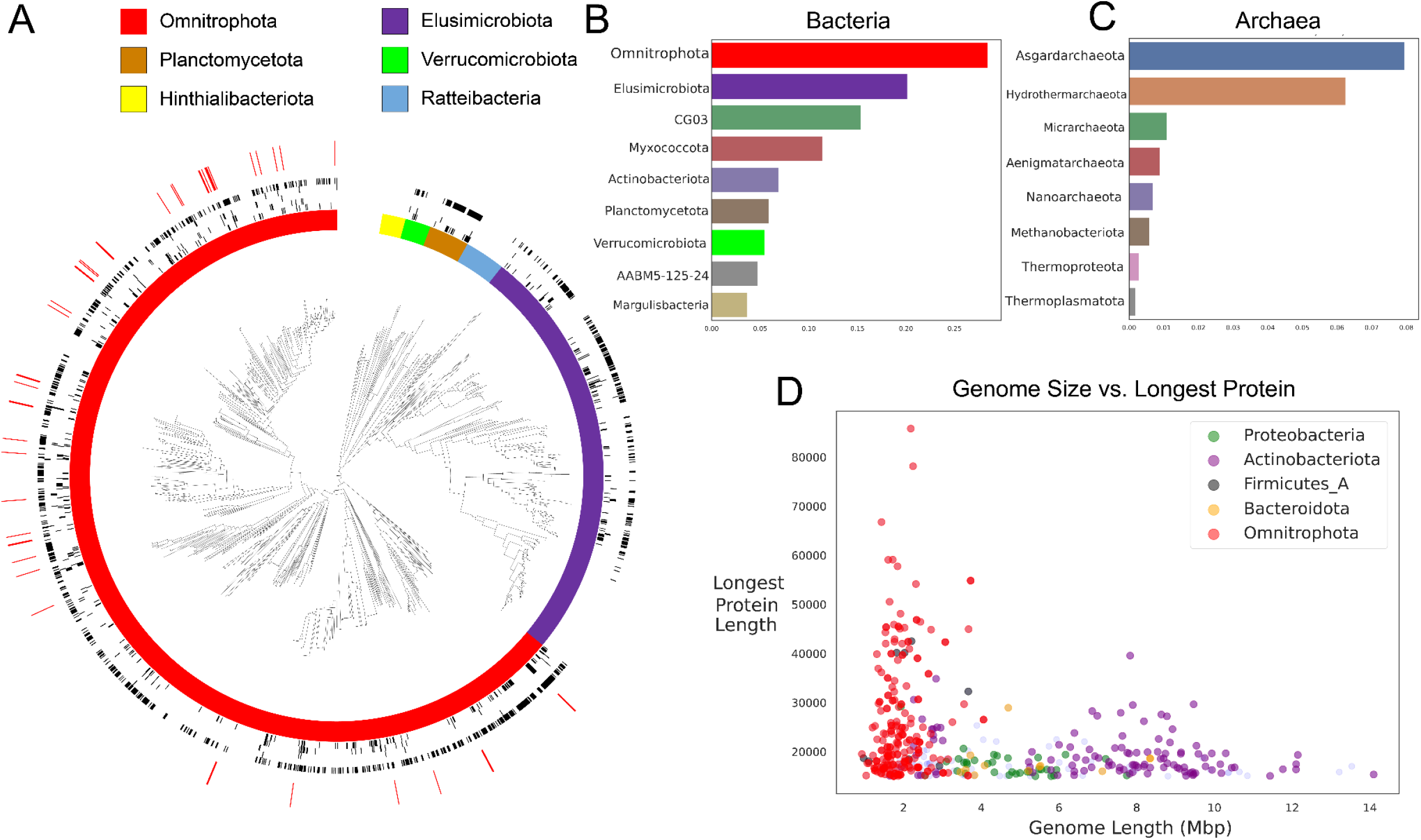
**A.** Phylogeny of representative phyla from the PVC supergroup: *Omnitrophota*, *Ratteibacteria*, *Verrucomicrobia*, *Hinthialimicrobiota*, *Planctomycetota* and *Elusimicrobiota,* constructed using 16 concatenated ribosomal proteins. Outer rings indicate the presence of proteins of a given length in a given genome; red lines indicate proteins of > 30,000 aa. Black ring annotations, from inner to outer, indicate proteins of: 5-10,000 aa in length; 10-15,000 aa; 15-20,000 aa; 20-25,000 aa; 25-30,000 aa. Red decorations on the exterior ring indicate presence of a predicted protein greater than or equal to 30,000 aa in length. **B.** Average copy number of predicted proteins > 10,000 aa among bacterial phyla with at least 10 representative genomes in the GTDB database. **C**. Average copy number of predicted proteins greater than 10,000aa among bacterial phyla with at least 10 representative genomes in the GTDB database. **D**. Genome size vs. length of the longest predicted protein in each genome for 5 phyla with highest total number of predicted protein sequences of > 10,000 aa. Note the high prevalence of proteins of >30,000 in length, despite relatively small genomes.

**Fig. 2.**
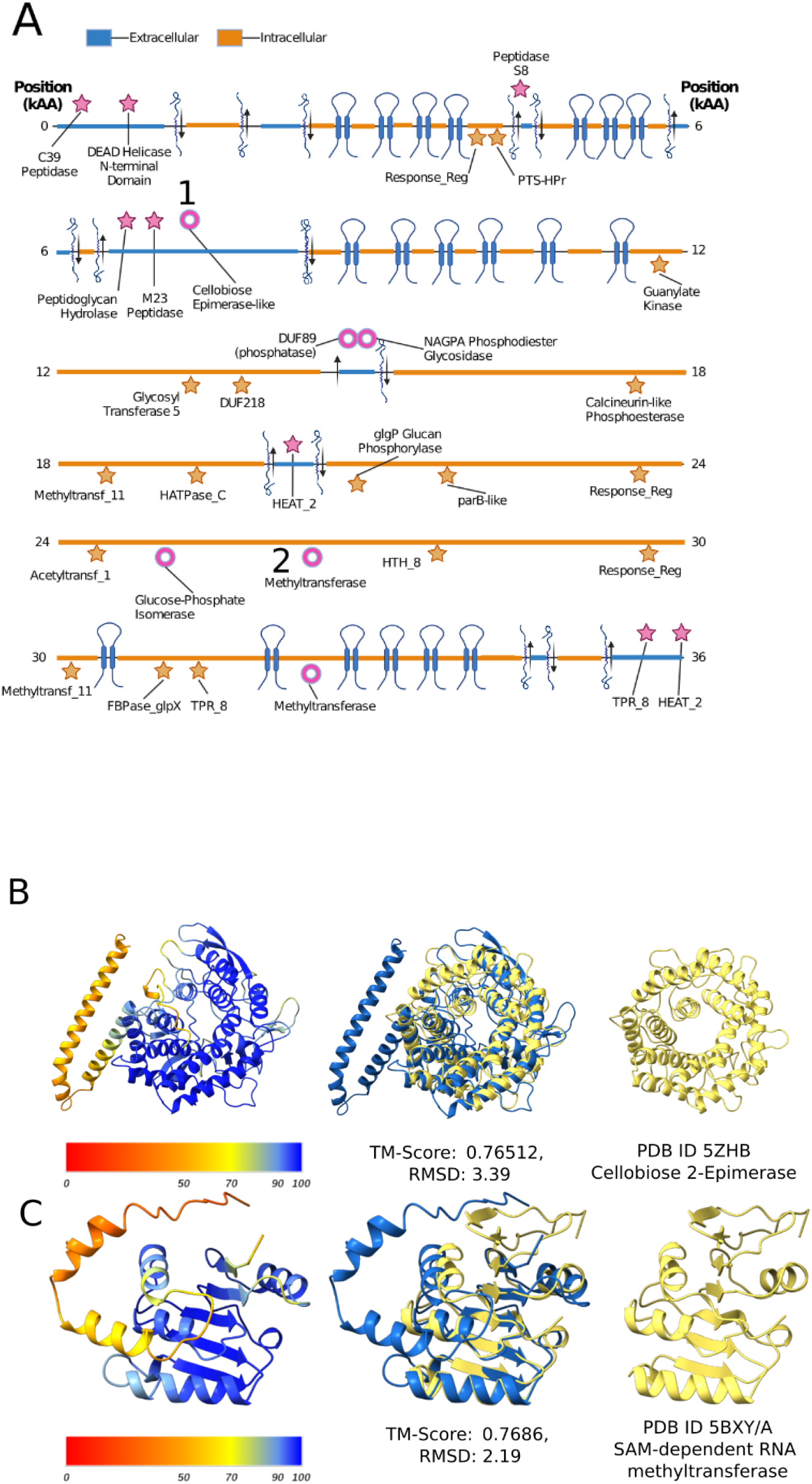
**A.** LC_01_metaspades_scaffold_444_73 domain diagram showing predicted extracellular localization and transmembrane helices. Pink stars indicate domains identified through HMM-based approaches; pink donuts indicate domains identified only through *in silico* structural analysis. **B**. *In silico* identified predicted domain structure homologous to reference Cellobiose-2-Epimerase (PDB 5ZHB). **C**. *In silico* identified predicted domain structure homologous to SAM-dependent RNA methyltransferase (PDB 5BXY).

**Fig. 3.**
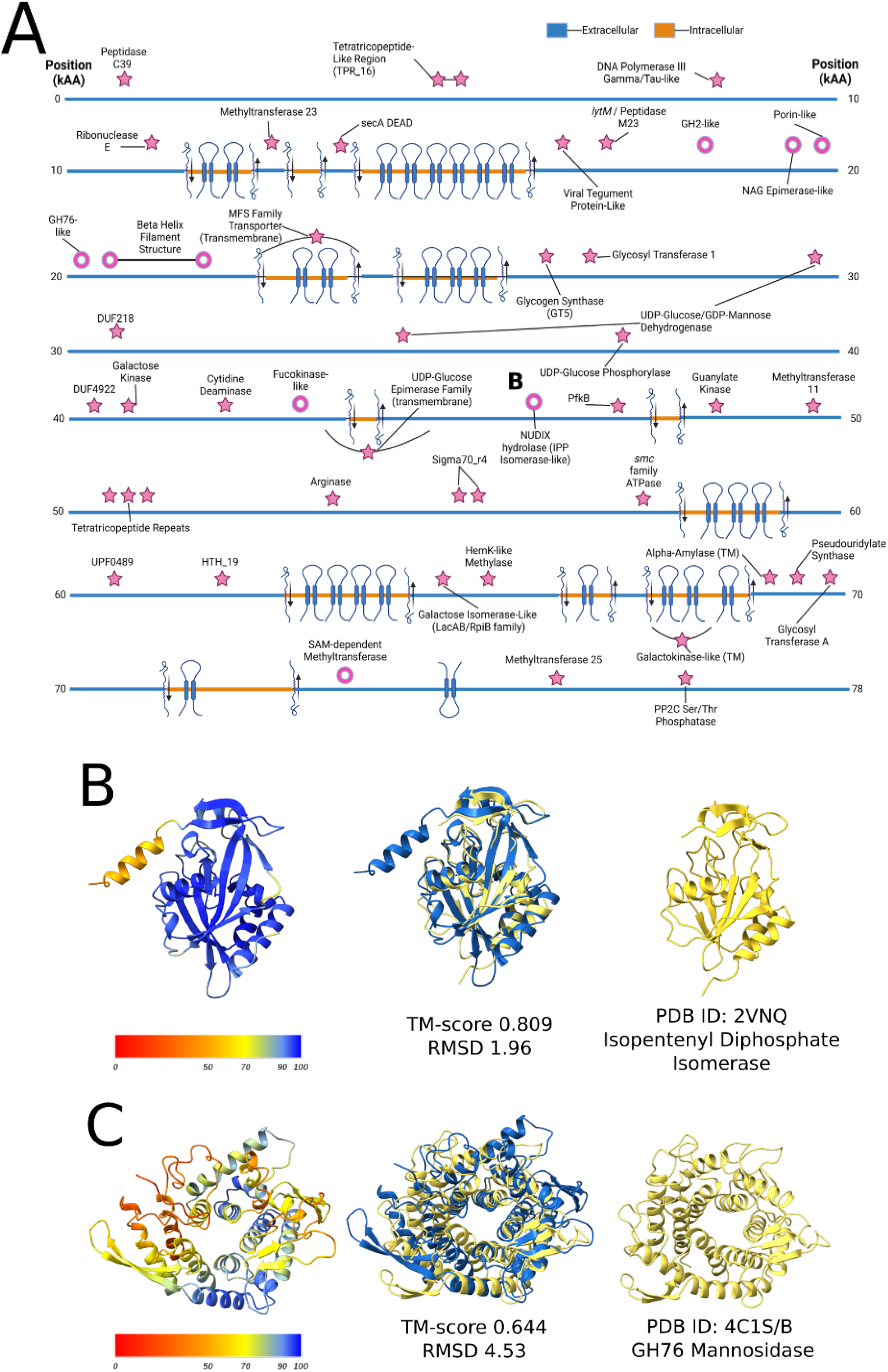
**A.** Domain diagram for Archaeovorin-type giant protein SR-VP_9_9_2021_34_2B_1_4m_PACBIO-HIFI_HIFIASM-META_416_C_1924. Pink stars indicate domains identified through HMM-based approaches; pink donuts indicate domains identified only through *in silico* structural analysis. **B.** *In silico* identified structure homologous to Isopentenyl Disphosphate Delta-Isomerase (PDB 2VNQ). **C**. *In silico* identified structure homologous to glycoside hydrolase family 76, which includes Mannosidase (PFAM 4C1S/B).

### New giant Omnitrophota proteins

To further investigate giant *Omnitrophota* proteins, we searched public databases for predicted protein sequences of ≥ 30,000 aa in length and identified them in genomes from a wide diversity of environment types (Methods). We further expanded this dataset by analysis of metagenomic datasets that we generated from a wetland located in Lake County, CA, a wetland and groundwater from Gunnison County, CO, circumneutral mine drainage in Napa County, CA, soil from a wetland and deep subsurface groundwater from sedimentary rocks from Japan. The wetland soils, mine drainage and deep groundwater all contain abundant Archaea that coexist with *Omnitrophota*.

Given the tendency of short read assemblies to generate fragments that do not capture entire giant genes, we obtained Nanopore and PacBio as well as Illumina datasets from two samples to enable manual curation of genes and genomes to completion. The long-read assemblies generated many long and some potentially complete genomes, but contain insertions/deletions of single bases that throw genes out of frame, prematurely terminating the giant genes. These errors were corrected using the Illumina reads. Errors also occur in the Illumina-only assemblies, which comprise the majority of the dataset. For example, in one instance, two predicted giant protein genes were located next to one another on the same contig. Correction of a local assembly error resulted in a single gene encoding a protein of 74,983 aa. In other cases, fragmented genes were extended via curation.

To augment giant gene sequence information we curated three *Omnitrophota* genomes to completion to determine the location of the giant genes. These genomes were verified by aligning illumina reads to the curated genomes at 100% identity and 100% coverage. One PacBio long read-based draft genome, previously assembled in two pieces, was curated to completion after the segments were joined by curation using illumina reads. The other PacBio long read-based draft genome genome was circular after assembly, and was polished to completion using illumina reads generated from the same sample. The third was assembled in two fragments from PacBio data, and joined together using Illumina and Nanopore reads generated from samples obtained at the same depth. The final *Omnitrophota* genomes are 1.6, 1.6, and 2.1 Mbp in length respectively, and show patterns of GC skew and cumulative GC skew that are expected for complete bacterial genomes, with two replichores. Interestingly (and unlike the vast majority of bacterial genomes), each replichore has genes almost all encoded on one strand (**Supp. Figures S3, S4, S5**: Genome Diagrams). The genomes encode *dnaA* at the origin of replication. The single giant protein encoding gene in each genome occurs around 100 kbp from the predicted origin of replication.

We obtained several draft *Omnitrophota* genomes from a metagenome from a swamp in Gunnison County, CO. Although the genes encoded protein sequences of ≥ 10,000 aa, the genes were incomplete. These partial proteins have regions of high homology to other giant *Omnitrophota* proteins. Similar results were obtained from groundwater and bog sediment microbial communities from the East River watershed in Gunnison County, CO, and from groundwater from the deep subsurface in Japan.

### Giant Omnitrophota genes

In total, 46 non-redundant gene sequences of >90,000 bp were identified within 1873 *Omnitrophota* genomes. The longest open reading frame is 257,412 bp in length (85,804 aa) and is encoded in an *Omnitrophota* genome obtained from wastewater.

To investigate whether these giant genes are translated as predicted, we analyzed the proteomics data from Kizina et al. 2022, in which *Vampirococcus* sp. LiM was shown to express a giant protein to a high degree when colocalized with *Methanosaeta.* Published spectra were converted to sequences through *de novo* peptide sequencing using Casanovo <u>(Yilmaz et al. 2022)</u>. Peptide fragment alignments were filtered using a threshold of ≥ 80% sequence identity and ≥ 80% peptide fragment query coverage. Even with this low threshold, only 381 peptide fragment sequences of average length 7.4 amino acids could be aligned to the giant protein from *V. archaeovorus LiM*. This is lower than the reported number of total spectral counts for this protein published in Kizina et al. 2022. This indicates reduced throughput using a *de novo* peptide sequencing approach when compared to. Full proteomic characterization of a giant protein from *Omnitrophota* will likely require a targeted enrichment or isolation of a giant protein-bearing *Omnitrophota* and higher throughput proteomics analyses.

To begin to explore the possible functions of giant proteins, we investigated the cellular localization of regions that encode detected functional domains, and found that the vast majority are extracellular (**Supp. Fig S1**) (**Supp. Fig S2**). However, some predicted localizations seem highly unlikely, suggesting that the transmembrane domain predictions are at times unreliable. Computational constraints allowed some, but not all, giant protein sequences to be analyzed with deepTMHMM <u>(Hallgren et al. 2022)</u>. For these transmembrane domain predictions, both the placement and number of predicted transmembrane helices, and thus prediction of extracellular and intracellular localization, often differed from the output of TMHMM <u>(Krogh et al. 2001)</u>. Thus, the complete distribution of transmembrane helices remains ambiguous. Domains on the giant proteins are often highly divergent from, and show low sequence similarity to, other domains of the same type from across Bacteria (**Fig. 4B**). Overall, most giant protein sequences contain numerous transmembrane helix domains, suggesting an association with the cell membrane.

**Figure 4.**
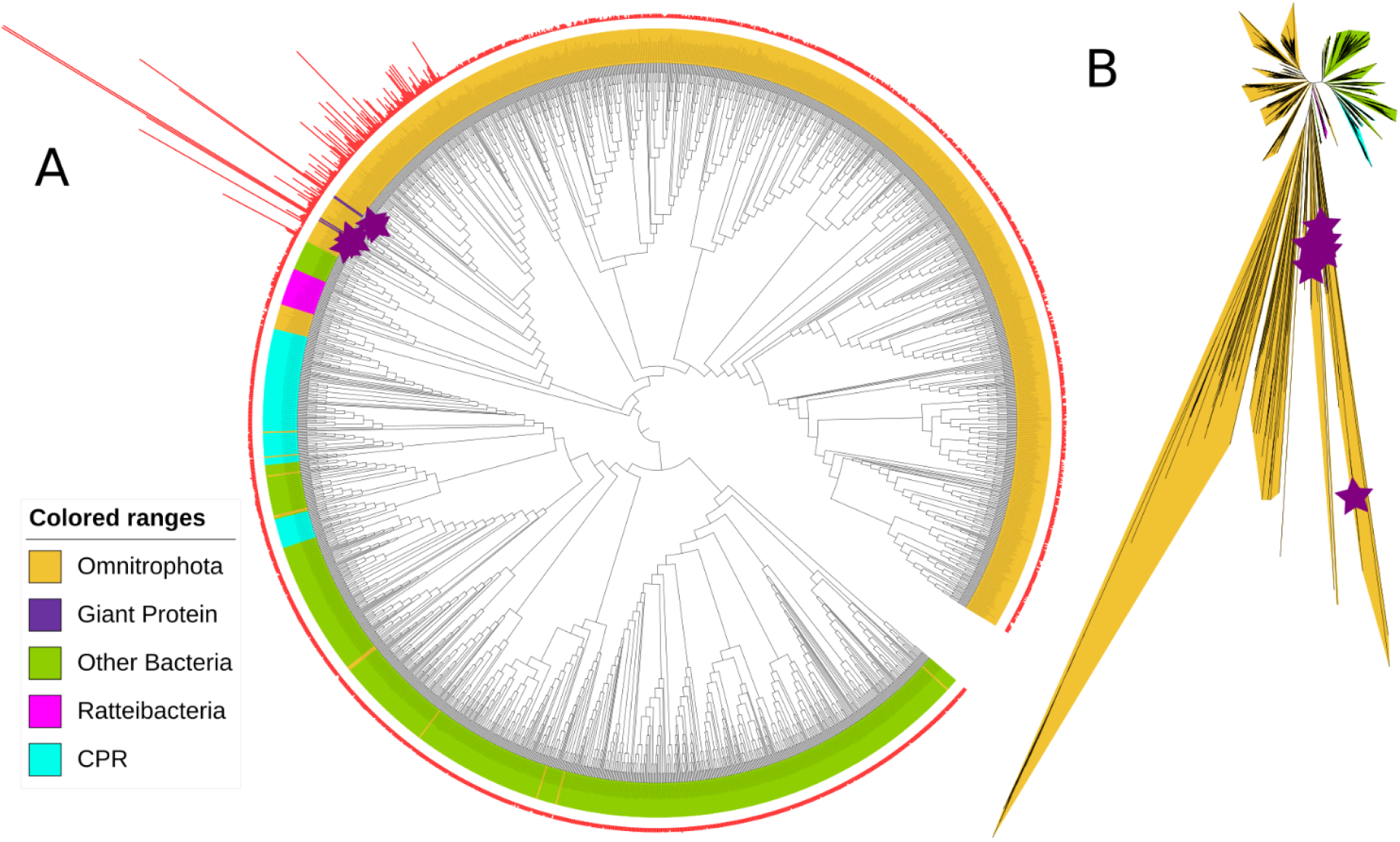
**A.** Cladogram of the SecA_DEAD domain across domain Bacteria, based on a maximum likelihood tree. Sequences obtained from giant predicted proteins are labeled with stars. Barchart on the outer ring indicates the size of the original ORF. **B.** Unrooted view of the phylogeny from subfigure A, visualizing branch lengths. Sequences obtained from giant predicted proteins are labeled with stars.

Full-length giant gene sequences universally contain predicted N-terminal regions that generally lack identifiable domains, with the exception of C39 peptidase domains near the N-terminus, which occur near predicted signal peptides. These domains also occur in bacteriocin and signal peptide preprocessing peptidases and as potential chaperones in type I secretion systems such as the RTX transporter HlyB <u>(Lecher et al. 2012)</u>. Others contain N-terminal domains with homology to peptidases of the S8 subtilisin family, known for autocatalytic protease activity <u>(Bratanis and Lood 2019) (Ikemura and Inouye 1988)</u>. The C39 and S8 peptidase domains might cleave the giant proteins into multiple smaller polypeptides. However, alternatively they may cleave other protein substrates.

The presence of numerous transmembrane domains implies that some regions of the giant proteins are extracellular. This raised the question of how these extracellular and transmembrane regions are secreted across and into the cell membrane. Two domains suggestive of involvement in protein secretion are common on predicted giant proteins of all subtypes. These are the overlapping SecA DEAD helicase and SecA preprotein binding domains that occur in the canonical SecA protein. Interestingly, they lack the scaffold and wing domain that comprises the membrane anchor of canonical SecA, raising a question of how these secretion components associate with the cell membrane <u>(Hunt et al. 2002)</u>. The adjacent transmembrane regions on the giant protein may provide a potential mechanism for membrane anchoring. SecA helicase domains from giant proteins form a phylogenetic clade distinct from other *Omnitrophota* SecA helicase domains and domains from outside the phylum. (**Fig. 5A** and **B**) and display exceedingly long branch lengths (**Fig. 5B**) suggestive of rapid evolution.

**Fig. 5:**
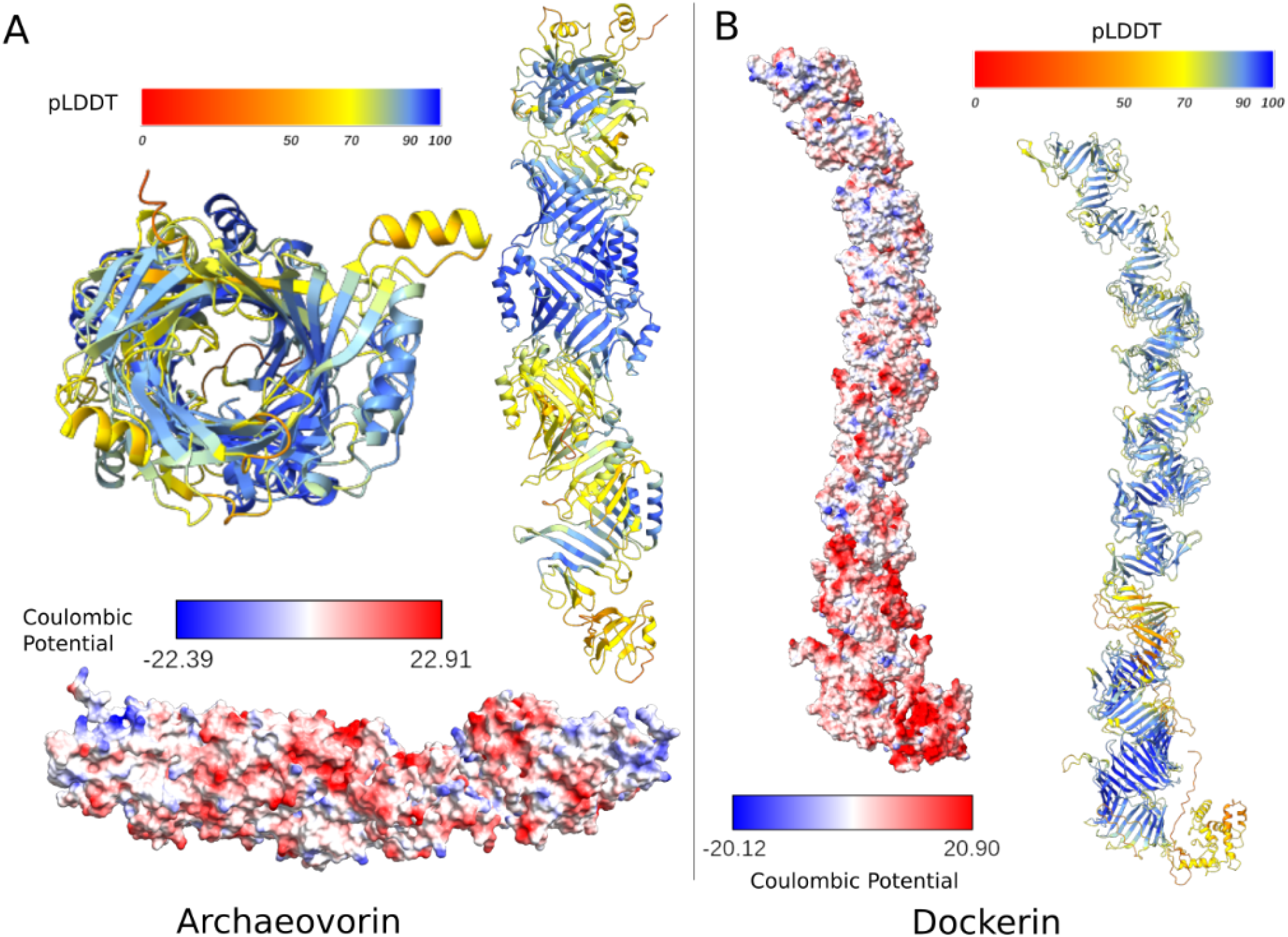
Large antiparallel beta-helix structure found in both main subtypes of *Omnitrophota* giant proteins. **A**) Antiparallel beta helix structure predicted from Archaeovorin-type protein occurring within the genome SR-VP_9_9_2021_34_2B_1_4m_PACBIO-HIFI_HIFIASM-META_416_C, showing external paired alpha helices. **B)** Antiparallel beta helix structure predicted from Dockerin-type sequence CAILPI010000004_73 showing external paired antiparallel beta sheets in place of paired alpha helices as observed in **A**.

Giant proteins containing SecA helicase domains are observed in the same complete genomes as normal-length SecA proteins that phylogenetically cluster separately. The *Omnitrophota* genomes also encode the SecYEG transmembrane channel, SecB chaperone, and SecD (that assists in translocation), although they are not syntenic with one another or with the giant genes. These, along with SecA (the ratchet protein), are the expected components of canonical Sec transport systems. In combination, these observations suggest that the SecA-like domains of *Omnitrophota* giant genes play a role that is distinct from that of canonical SecA and are potentially involved in the export process of the giant protein itself.

The sets of predicted domains of the 46 giant *Omnitrophota* proteins suggest their separation into two categories. The first category consistently contains glycosyl transferases, methyltransferases, glycosyl hydrolases, peptidases, SAM-dependent methylases, and NUDIX hydrolases. These domains are also present in the giant protein from *Velaminicoccus archaeovorus* (**Supp. Fig S2**), which predate archaea. Thus, we refer to this type of *Omnitrophota* giant proteins as Archaeovorins and nominate this *V. archaeovorus* protein to be the type sequence. The second and more rarely observed category features a large number of Dockerin and Thrombospondin type 3 repeats. Dockerins are cohesion components of extracellular complexes and Thrombospondin type 3 repeats are conserved regions of extracellular proteins that regulate cell-to-matrix interactions <u>(Pagès et al. 1997; Kvansakul et al. 2004)</u>. We refer to these predicted membrane-anchored extracellular proteins as Dockerin-type giant proteins.

### Archaeovorins

The most frequently observed domains in Archaeovorins are HEAT repeats that are exclusively detected in predicted extracellular regions of Archaeovorin protein sequences (**Fig. S1)**. These domains are crucial for assembly of multi-enzyme complexes and could play this role in *Omnitrophota,* or serve as protein-binding surface antigens. The best studied examples are in the cellulosome <u>(Carvalho et al. 2003)</u> where they form cohesin domains <u>(Neuwald and Hirano 2000)</u>.

The second most common domains in Archaeovorins are glycosyl transferases (GT). Most prevalent are GT family 5 domains with homology to glycogen synthase domains that could be involved in construction or breakdown of a branched glycopolymer. Also abundant are GT 1 domains, which are involved in various polysaccharide transformations. While their roles remain uncertain, they could potentially be involved in converting cell wall degradation products to carbon storage compounds (e.g., glycogen), glycosylation of extracellular proteins, or removal of glycosyl moieties from glycoproteins.

Archaeovorins often feature tetratricopeptide repeat (TPR) superfamily domains that bind N-acetyl glucosamine-containing substrates such as cell wall compounds including peptidoglycan or glycoproteins <u>(Iyer and Hart 2003)</u>. They are also involved in protein-protein interactions <u>(Cerveny et al. 2013)</u>. Both GT 1 and TPR repeat domains co-occur in UDP-N-acetylglucosamine 4-epimerase, an enzyme that interconverts UDP-N-acetyl-D-galactosamine and UDP-N-acetyl-D-glucosamine, possibly enabling the galactose-rich products of glycoprotein degradation to be fed into glycolysis. This domain combination occurs in many other proteins in public databases, but unfortunately their functions are unknown. Although many other Archaeovorin domains were commonly and confidently identified, we are unable to infer potential roles (**Figure S1**).

A less common domain identified in Archaeovorins is also found in SLT lytic transglycosylases that break the beta-1,4 glycosidic bond of murein. This may indicate roles for Archaeovorin protein regions in binding to and degrading cell wall compounds such as murein or pseudomurein. Given that pseudomurein occurs in archaeal cell walls, we speculate that certain *Omnitrophota* use Archaeovorins to attach to and degrade *Methanosarcinales* cell walls.

Less common, but perhaps easier to assign roles for, are domains homologous to those of triosephosphate isomerase, glucose-phosphate isomerase, fructose 1,6-bisphosphatase, phosphoglucomutase/phosphomannomutase, and fructose-bisphosphate aldolase. These are predicted to be in an extracellular region, but it is quite possible that an overlooked transmembrane resulted in the wrong localization. If extracellular, these domains may enable pre-processing of sugars from the extracellular environment prior to import into the cytoplasm for further consumption via glycolysis. Archaeovorins also frequently contain pyruvate-diphosphate kinase domains that interconvert pyruvate and phosphoenolpyruvate.

### Dockerin-type giant proteins

The second category of giant genes encode dockerin domains that do not occur in Archaeovorins. There are just three examples of these proteins with ≥ 30,000 aa. Dockerin-type giant proteins contain numerous repetitive domains, including EF-hand repeats (of which Dockerin repeats are comprised) and bacterial thrombospondin type 3 repeats. All three proteins are likely incomplete due to unresolvable assembly fragmentation due to the repeats. Notably, both Dockerin domains and type 3 thrombospondin domains contain calcium binding sites <u>(Kvansakul et al. 2004)</u> and both display homology to component domains of the cellulosome that are involved in attachment <u>(Dai et al. 2017) (Rincon et al. 2003)</u>. They also contain discoidin repeats (PFAM F5_F8_type_C) that occur in N-acetylgalactosamine binding endoglucanases <u>(Mathieu et al. 2010)</u>.

Dockerin type giant proteins lack predicted transmembrane domains along the majority of the N-terminal portion of the gene. Those with complete C-termini show predicted transmembrane and intracellular regions. Given their predicted functional domains, we suggest that the Dockerin-type giant proteins are involved in attachment, possibly to a host or prey cell surface.

Numerous domains found within Dockerin type giant proteins are suggestive of binding to and degradation of glycosyl moieties, such as DUF1080 domains, which occur in proteins with beta-glucosidase and beta-mannosidase activity <u>(Elcheninov et al. 2017)</u> and the ILEI domain, which has been noted for its homology to O-linked mannose 1,2-N-acetylglucosaminyltransferase <u>(Kuwabara et al. 2016)</u>. GNAT family acetyltransferases, also observed on Archaeovorins, are common on Dockerin type giant proteins, and may be involved in deacetylation of cell wall compounds such as GlcNAc and Glucuronate. Thus, like Archaeovorins, an important role for Dockerin-type giant genes may be in processing of cell surface polymers.

One Dockerin-type sequence contains C-terminal C39 peptidase-like domains, also observed in Archaeovorins, and similarly these may be involved in proteolytic cleavage of a larger propeptide. Transmembrane anchor domains exist near the C-terminus, followed by secA related domains, as seen in Archaeovorins. Similar to the SecA helicase domains in Archaeovorin sequences, these SecA helicase domains may play a role in secretion of the giant protein itself, or of accessory proteins. Overall, the presence of domains predicted to be involved in transferring of acetyl groups, protein secretion, and proteolytic processing of larger proproteins in both Dockerins and Archaeovorins indicates some high level similarity between these giant proteins.

### *In silico* protein structure prediction of unannotated sequence

As large stretches of the predicted giant proteins lack identifiable functional domains, we explored the utility of *in silico* structural prediction software to uncover regions with tertiary structure that may provide functional clues (**Fig. 4**). However, it is infeasible to predict structures for proteins of >30,000 amino acids. It is possible that giant proteins could be cleaved (informatically) into many independent polypeptides for which predictions would be practical. However, the absence of information about cleavage points precluded this approach. Thus, we developed a new approach for detection of domains in giant proteins using *in silico* structural biology methods.

As structural prediction is difficult for proteins with numerous transmembrane domains, we subdivided the giant proteins in ways that excluded regions with transmembrane domains. Regions for which functional domains had not been identified were subdivided into partially redundant, overlapping 1000 aa fragments with a step size of 500 aa and their predicted folded structures calculated using Alphafold2 (see Methods). Overall, we found that reducing the sequence length was very important for identification of putative domains in giant proteins. This resulted in the prediction of several large (>1000 aa) regions of continuous folded tertiary structure on both Archaeovorin- and Dockerin-type sequences. In some cases, predicted tertiary structure extends across multiple 1000 aa fragments (**Fig. 5**).

Both Archaeovorin- and Dockerin-type protein regions are predicted to assemble into remarkable ∼1500 aa tubule-like structures comprised of a long, single-stranded antiparallel helix comprised of antiparallel beta sheets with regularly interspaced exterior alpha helices and a hollow interior (**Fig. 5**). This antiparallel beta helix-based structure is novel, but shares some features with RHS repeat proteins including type VI secretion system (T6SS) and ABC toxin complex effector proteins that also have exterior antiparallel beta strands that form a helical structure <u>(Meshcheryakov et al. 2019)</u>. Additionally, our novel helical structure contains some similarity to the VgrG spike of the *E. coli* type VI secretion system. Although the spike protein has a similar internal radius it has only a single ring of paired external alpha helices and contains a parallel rather than an antiparallel beta sheet structure <u>(Flaugnatti et al. 2020)</u>. Thus, to our knowledge, the giant hollow protein protrusions are based on an unusual fold that forms a novel structure.

*In silico* protein structure prediction revealed a region folded with high confidence in the middle of an extracellular segment of a Dockerin-type protein. The predicted structure contains many unannotated repeating domains that form antiparallel beta barrels of ∼200 amino acids in length that are linked by alpha helices. The beta barrel region shows highest homology to carbohydrate-binding modules (CBMs) of family 35, which are known to bind specifically to cell wall-associated glucuronate and galacturonate-containing compounds <u>(Montanier et al. 2009)</u>. However, CBM family 35 lacks the linking alpha helices. The alpha helices align and show high structural homology to canonical dockerin domain-containing proteins that form complexes with cohesin proteins (**Fig. 6D**). In canonical dockerin domain-containing proteins, the alpha helices are formed by a single polypeptide but here, the alpha helices flank the CBM family 35-like beta barrel. (**Fig. 6C,D**). These observations suggest the existence of a fold that is a natural combination of a carbohydrate binding module and a dockerin. Regions containing this repeated domain show ladder-like macrostructure.(**Fig. 6A**)

**Fig. 6.**
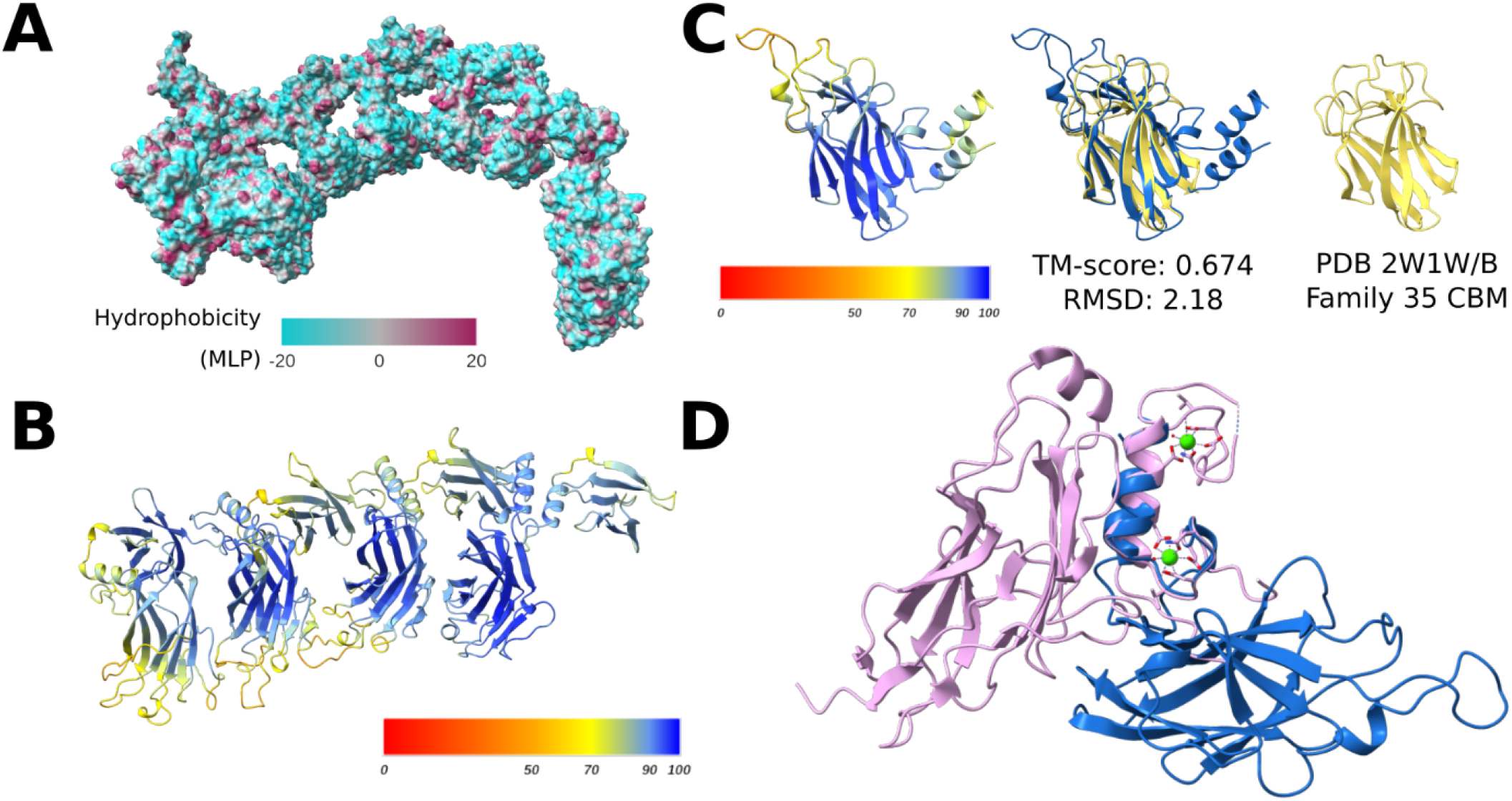
Unannotated region of Dockerin-type protein CAILPI010000004_73 containing Dockerin/Cohesin-like folded subunits. **A.** Hydrophobicity surface of 2500 aa subregion containing repeating dockerin/CBM35 complex-like subunits showing predicted higher-order tertiary structure. **B.** Independently folded 1000 aa subfragment of the previous structure showing alternating beta barrels and twinned alpha helices. Color indicates confidence in folded structure as measured by pLDDT. **C**. Individual subunit of repeating structure shown in **B** aligned to highest scoring PDB hit, a carbohydrate binding module of family 35. Color bar shows pLDDT. **D**. Canonical dockerin/cohesin pair (PDB 2y3n A,B: pink, left) and repeating subunit from Dockerin-type giant protein (blue, right). Alignment was performed between the Dockerin chain and alpha helical region of the giant protein sequence. Green indicates calcium ligands bound to PDB reference.

**Fig. 7:**
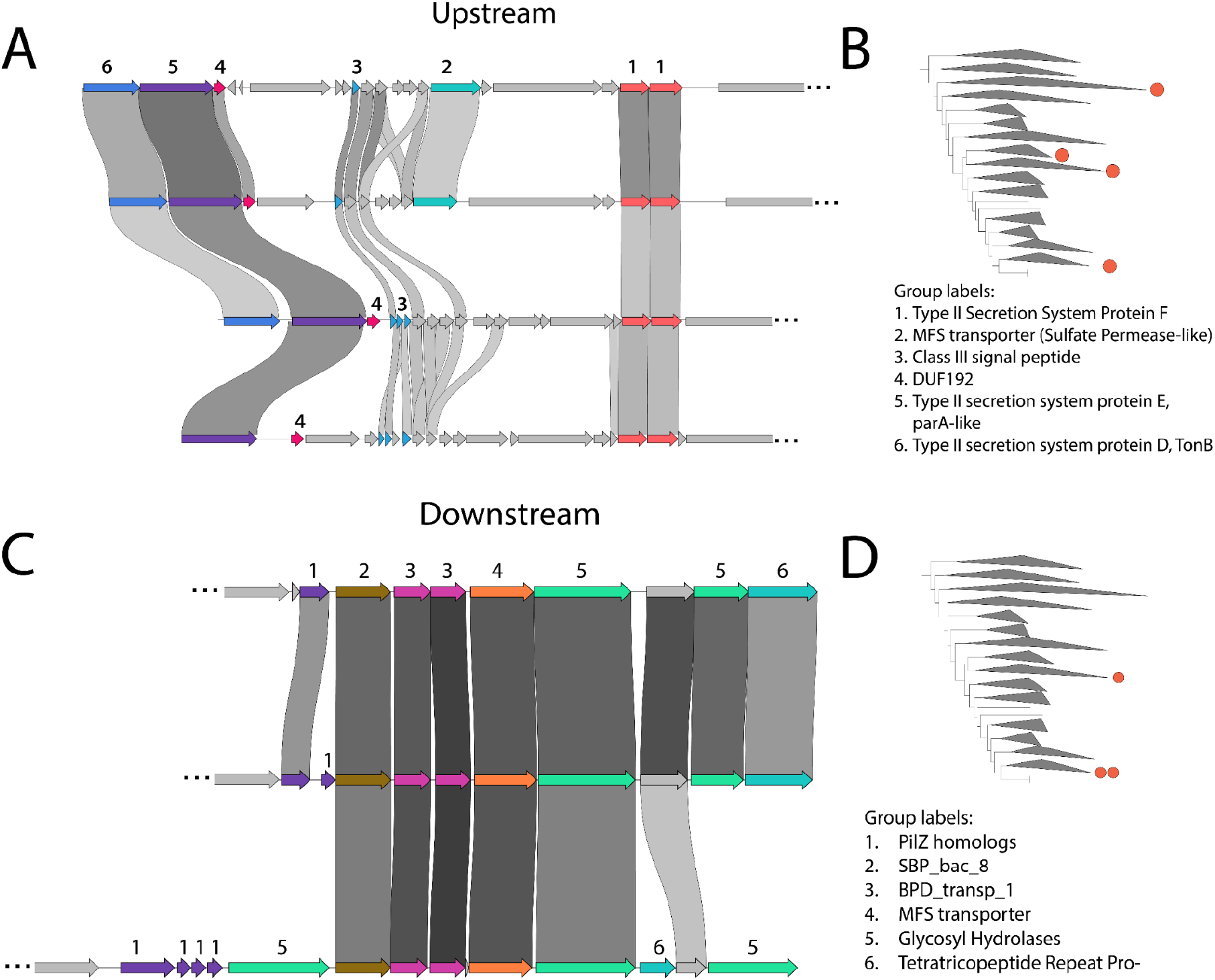
**A.** Conserved type II secretion system-like locus upstream of giant gene sequences. Links indicate pairwise amino acid identity of at least 30%. **B.** Collapsed *Omnitrophota* ribosomal phylogeny, as displayed in Figure 1, with red dots indicating phylogenetic positions of genomes containing genomic loci displayed in subfigure **A. C.** Conserved *pilZ* genes and soluble binding protein-dependent carbohydrate transport systems downstream of giant genes. **D.** Collapsed *Omnitrophota* ribosomal phylogeny with red dots indicating phylogenetic positions of genomes containing genomic loci displayed in subfigure **C.**

### Conserved functionalities encoded in proximity to giant genes

The groups of proteins encoded prior to and following the *Omnitrophota* giant genes are relatively conserved. The giant proteins, regardless of differences in domain content, are generally preceded by a secretion system with more than 10 subunits, many of which display homology to proteins from type II and type IV secretion systems, including the type II secretion system proteins *gspD*, *gspE* and *gspF*, as well as pilin formation systems encoded by the *rcpC* genes and a conserved peptidyl-prolyl cis-trans isomerase containing a Rotamase_2 domain (**Fig. 5A**). Additionally, many small genes occur nearby with homology to class III signal peptides. Although other secretion system proteins in these regions are less well conserved, this secretion system locus is conserved across a wide range of *Omnitrophota* genomes from across the phylum (**Fig. 5B**).

The majority of giant genes are immediately followed downstream by one or multiple small *pilZ* homologs, which bind cyclic di-GMP and could interact with the GGDEF diguanylate cyclase domains observed in many Archaeovorin sequences. Further downstream of the majority of giant genes are numerous extracellular solute binding proteins, binding protein-dependent transporters, and glycosyl hydrolases. Some genes contain the sbp_bac_3 domain, which is shared with membrane-bound lytic murein transglycosylases. The function of these genes is consistent with the hypothesis that they, along with the nearby giant genes, are involved in prey cell wall binding and degradation.

### Comparison with giant bacterial proteins outside the phylum *Omnitrophota*

The most frequently observed giant genes outside of the *Omnitrophota* originate in genomes from *Actinobacteriota*. Some giant actinobacterial proteins contain secA preprotein translocase and SecA_DEAD helicase domain but lack the scaffold and wing subunit, as found in giant genes from *Omnitrophota*. Other domains that are shared by giant proteins in *Actinobacteria* and *Omnitrophota* include the GGDEF diguanylate cyclase domain, serine/threonine phosphatases, CheY-like receiver domains, NUDIX hydrolases, and LPXTG-anchored adhesin domains. However, these proteins contain far fewer detectable domains on average than similarly sized proteins from *Omnitrophota*, but this could be due to problems with domain identification, as encountered in *Omnitrophota* giant proteins.

<u>One example</u> of a giant gene, which codes for a protein 36,805aa in length, occurs in *Chlorobi* from the consortium named *Chlorochromaticum aggregatum*, in which a *Chlorobi* episymbiont attaches to the surface of *Symbiobacter* spp. cells <u>(Frostl and Overmann 1998)</u>. Domains identified in this protein include repeat sequences often observed in giant proteins, domains involved in attachment, toxin-related domains, and an M36 family metallopeptidase domain often involved in cleavage of cell surface proteins by pathogenic fungi <u>(Rawlings et al. 2018)</u>. This genome also encodes a 25,000 aa protein that features fibronectin and haemagglutinin-like repeats, which have been previously hypothesized to play a role in host adhesion <u>(Liu et al. 2013)</u>.

A unique class of giant genes was observed in the class *Clostridia*, family *Oscillospiraceae*. The predicted proteins are comprised entirely of repetitive protein domains, primarily RCC1 (PFAM ID PF00415) <u>(Swart et al. 2020) (Stevens and Paoli 2008)</u>. RCC1 domains fold into seven-bladed □-propeller structures, which are understood to evolve through amplification and subsequent diversification within individual genes <u>(Pereira and Lupas 2022)</u>. In contrast to those in *Omnitrophota* and in *Archaea*, *Oscillospiraceae* RCC1-type giant genes have a single repeating protein domain along the entire length of the sequence, similar to *swmB* and Titin. *Oscillospiraceae* RCC1-type giant genes were primarily found in gut metagenomes from humans, mice, and rabbits, although one was observed in a genome of the genus *Pseudoflavonifractor* in an anaerobic chlordecone-degrading enrichment from soil <u>(Chaussonnerie et al. 2016)</u>. These proteins lack transmembrane anchor regions and are predicted to localize to the extracellular. Bacterial RCC1 proteins are known to play roles in pathogenicity. Some bacterial RCC1 proteins from *Legionella* are secreted into eukaryotic cells and promote growth of *Legionella* within the host cell <u>(Swart et al. 2020)</u>. An RCC1-like beta propeller has been observed within an insecticidal toxin complex from *Yersinia entemophaga* tha talso bears Rhs repeats which adopt similar structures to the beta helix filaments from *Omnitrophota* giant proteins <u>(Hurst et al. 2011)</u>.

Other large genes from class *Clostridia* are between 10,000 and 30,000 aa in length, similarly obtained from mammalian gut metagenomes <u>(Mitchell et al. 2018)</u>, These proteins contain numerous other beta propeller-forming domains such as the Listeria-Bacteroides repeat domain and YG repeat glucan-binding domain, potentially indicating a function involved in binding to surface glycoproteins of a host or prey cell <u>(Haas and Banas 2000)</u>. Additional protein domains observed on *Clostridia* giant predicted proteins include Dockerin repeats, which are featured prominently in giant proteins from *Omnitrophota*, as well as surface antigen-associated domains such as Leucine-rich repeats and fibronectin-binding autotransporter adhesin ShdA <u>(Plaza Onate et al. 2022)</u>. From these observations it appears that giant proteins, both in *Omnitrophota* and in other bacteria, may often play roles in adhesion to the surfaces of other cells. Overall, giant proteins from *Omnitrophota* and large proteins from these bacteria of these three phyla appear to share functional similarities and potentially all play roles in bacterial predation.

## Conclusions

The existence of giant proteins, sometimes exceeding 85,000 amino acids in length, challenges our understanding of protein size limits in microorganisms. An important question raised by their presence is whether the transcript and/or translated products function as one sequence or are cleaved into smaller transcripts or multiple peptides. The domains homologous to bacteriocin preprocessing endopeptidases as well as subtilisin family proteases suggest autocatalytic proteolysis into smaller polypeptide products. Regardless of if, and at what stage, cleavage occurs, this study sheds light on an interesting biological strategy apparently confined to a subset of bacterial lineages and underscores the vast, unexplored genetic potential hidden within uncultivated microbial communities.

Giant genes in the genomes of bacteria from the candidate phylum *Omnitrophota* are surprisingly common. They may be a conserved feature of these bacteria, but genome fragmentation precludes a full assessment of their ubiquity. Notably, giant genes are found on all circular *Omnitrophota* genomes generated to date. The origin and evolution of giant genes, including the incorporation of new domains and potential elongation by duplication of repetitive domains such as Dockerins and Thrombospondins, remains a topic with room for further exploration. Functional domains encoded within giant proteins are diverse, and suggestive of roles in cell surface attachment and cell surface polymer degradation. Future research could unravel the roles these giant proteins play in shaping microbial interactions and ecosystem dynamics.

The giant genes occur next to conserved upstream loci encoding secretion systems, and downstream from machinery for polysaccharide import to the cell. The gene content of these loci is consistent with potential roles in predation. Experimental evidence already confirms a predatory lifestyle for some *Omnitrophota* (Kizina et al. 2022), but biochemical evidence is needed to investigate the enzymatic functions of the domains and confirm the role of these genomic regions in predation.

Giant proteins predicted from genomes other than *Omnitrophota* often have domains with similar predicted functions to those from *Omnitrophota*, and in some cases share many domains with both Archaeovorin and Dockerin-type *Omnitrophota* giant proteins. These proteins often contain domains indicative of roles in cell-cell adhesion, as well as SecA-like domains similar to those found on *Omnitrophota* giant proteins.

The method we developed to enable *in silico* protein structural prediction of unannotated regions within the *Omnitrophota* giant proteins revealed several unusual high-confidence folded structures. For example, repeating subunits composed of a carbohydrate binding module-like fold and a dockerin-like fold form an elongate, ladder-like structure. The method also brought to light novel domains that form highly symmetric beta helices. The resulting tubule-like structure was predicted in both Archaeovorin and Dockerin type giant proteins, suggesting a potentially conserved structure across multiple subtypes of giant proteins.

The discovery of genes, particularly in a phylum known for ultra-small genome and cell sizes, raises intriguing questions about the pressures and ecological niches that motivate the evolution of very large open reading frames.

## Methods

### Discussion of Proprietary Data Sources SRVP

#### Sampling and nucleic acid extraction

Wetland soils were meticulously sampled during the late autumn months of October and November in the years 2017, 2018, 2019, 2020, and 2021 for the purpose of investigating microbial communities. Notably, the DNA extraction process involved gentle mixing in a temperature-controlled water bath at 65°C, as opposed to vigorous vortexing, to minimize potential degradation. This purification process included an overnight precipitation step at −20°C, followed by centrifugation at room temperature for 30 minutes. A secondary purification step involved a cold 75% Ethanol wash and another 30-minute centrifugation at room temperature. Following these steps, the DNA pellet was expertly resuspended in the provided elution buffer from the Qiagen kit. DNA was extracted from each sample (DNeasy PowerSoil Pro) and submitted for Illumina sequencing (150-bp or 250-bp reads) at the QB3 facility, University of California, Berkeley. Illumina reads were adapter and quality trimmed using sickle. Filtered reads were assembled using IDBA-UD <u>(Peng et al. 2012)</u>and MEGAHIT <u>(Li et al. 2015)</u>. Pacbio reads were assembled using hifiasm-meta <u>(Feng et al. 2022)</u>. Nanopore data was generated as described previously <u>(Schoelmerich et al. 2023)</u>. Metagenomic binning was performed using VAMB <u>(Nissen et al. 2021)</u>, MetaBAT2 <u>(Kang et al. 2019)</u> and MaxBin2 <u>(Wu et al. 2016)</u> in combination with manual binning performed on ggKbase. Gene predictions were established using Prodigal <u>(Hyatt et al. 2010)</u>.

### Corona Mine

Two metagenomic samples were taken from circumneutral (pH 6.1), Fe-rich mine drainage at the Oat Hill Mine, Napa County, CA, USA. These samples were size fractionated in order to capture ultramicroscopic microorganisms, using a 2.5 micron and 0.1 micron filter in sequence, and *Omnitrophota* genomes were concentrated in the 0.1 micron fraction. DNA was extracted using the Qiagen PowerSoil kit. Sequencing was done in two runs for each of the 0.1 and 2.5 micron filter samples; once with 150bp paired-end illumina reads, and once with 250bp paired-end illumina reads.

Metagenomic assembly was performed using metaspades.py with default parameters. Coverage of resulting metagenomic contigs was assessed by aligning reads using bbmap.sh with the following parameters: minid=0.96 idfilter=0.97 ambiguous=random. Metagenomic binning was performed using VAMB <u>(Nissen et al. 2021)</u>, MetaBAT2 <u>(Kang et al. 2019)</u> and MaxBin2 <u>(Wu et al. 2016)</u> in combination with manual binning performed on ggKbase. Resulting bins were consolidated with DAS Tool <u>(Sieber et al. 2018)</u> and further refined via manual inspection and curation on ggKbase. Genomes obtained from these metagenomes which were included in this analysis are located in **Supplementary Data S1**.

### Gossfen

One metagenomic sample was obtained from a bog site in the East River watershed, Gunnison County, CO, USA. DNA was extracted using the Qiagen PowerSoil kit and sequenced with 150bp paired-end illumina reads. Metagenomic assembly was performed using metaspades.py with default parameters. Coverage of resulting metagenomic contigs was assessed by aligning reads with bbmap.sh with the following parameters: minid=0.96 idfilter=0.97 ambiguous=random. Metagenomic binning was performed using MetaBAT2, MaxBin2, and manual binning on ggKbase. Resulting bins were consolidated with DAS Tool and further refined via manual inspection and curation on ggKbase. Of the 81 bins recovered from this sample, 43 were labeled as near complete based on the presence of single copy genes and ribosomal proteins <u>(Sieber et al. 2018)</u>. Five additional viral bins were recovered, three of which are megaphages over 300kbp in length. Near complete genomes from this sample will be submitted to NCBI upon publication.

### Shumway

One metagenomic sample was obtained from well water at a site in the East River watershed, Gunnison County, CO, USA. Water was extracted from a depth of 200 ft and then microbial communities were captured in bulk using a 0.1 micron filter. DNA was extracted using the Qiagen PowerSoil kit and sequenced with 150bp paired-end illumina reads. Metagenomic assembly was performed using IDBA_UD with default parameters. Binning was performed using MetaBat2, Maxbin2, and manual binning with ggKbase. Resulting bins were consolidated using DAS Tool and further curated via manual inspection on ggKbase. Of the 67 genomes obtained from this sample, 36 were labeled as near complete based on the presence of single copy genes and ribosomal proteins. Near complete genomes from this sample will be submitted to NCBI upon publication.

### Hyporheic

Multiple samples of hyporheic zone water were obtained from the East River, Gunnison County, CO, USA. Tubes were buried 30 cm below the riverbed and allowed to incubate for 24 hours before water was pumped out of this region and used to capture the microbial communities in bulk using a 0.1 micron filter. DNA was extracted using the Qiagen PowerSoil kit and sequenced with 150bp paired-end illumina reads. Metagenomic assembly was performed using metaspades.py with default parameters. Binning was performed using MetaBat2, Maxbin2, and manual binning with ggKbase. Resulting bins were consolidated using DAS Tool and further curated via manual inspection on ggKbase. Genomes obtained from these metagenomes which were included in this analysis are located in **Supplementary Data S1**; these bins will be submitted to NCBI GenBank upon publication.

### BJP

Multiple samples (or One sample) of deep groundwater were obtained from the Horonobe Underground Research Laboratory, Hokkaido, Japan. DNA was extracted using the Extrap Soil DNA Kit Plus ver. 2 (Nippon Steel and Sumikin Eco-Tech Corporation, Tsukuba, Japan) and sequenced with 150 bp paired-end Illumina reads. Metagenomic assembly was performed using *de novo* using IDBA-UD with the following parameters: -mink 40, -maxk 100, -step 20 and -pre correction. Bining was performed manually with ggKbase. Genomes obtained from these metagenomes which were included in this analysis are located in **Supplementary Data S1** and will be submitted to NCBI upon publication.

### Discussion of Public Data Sources

Genomes from the BV-BRC database taxonomically annotated as *candidatus Omnitrophica* were downloaded on May 18, 2023 using a custom python script utilizing the BV-BRC Command Line Interface, BVBRC_genome_grabber.py (**Supplementary Data S2, S3**). Genomes for *Elusimicrobia*, *Verrucomicrobia* and *Planctomycetes* were downloaded from BV-BRC using BVBRC_genome_grabber.py on June 9, 2023 by searching for genomes taxonomically annotated as ‘Elusimicrobia’, ‘Verrucomicrobia’, and ‘Planctomycetes’ respectively.

The 10,000 longest predicted protein sequences available on BV-BRC were downloaded on June 29, 2023 (**Supplementary Data S4**. Sequences containing gaps, represented by X characters in the predicted amino acid sequence, were removed, leaving 7,478 proteins for further analysis, with minimum length 10,624aa.

Reference proteins from UniProt <u>(UniProt Consortium 2023)</u> were obtained by searching for proteins with hits to the PFAM domains which were chosen for phylogenetic analysis. Only reviewed sequences with the domains of interest were downloaded and included in this analysis.

The public datasets in which giant proteins were detected were from landfill leachate metagenomes <u>(Grégoire et al. 2023)</u>, anaerobic digester sludge in the United States <u>(Mei and Liu 2022)</u> and China <u>(Liu et al. 2021)</u>, lake water in Spain <u>(Cabello-Yeves et al. 2022)</u> and Finland <u>(Buck et al. 2021)</u>, subsurface fracture fluids <u>(Casar et al. 2021)</u> in the United States, groundwater in New Zealand <u>(Mosley et al. 2022)</u> and Finland <u>(Bell et al. 2022)</u>, hydrocarbon field communities in Canada <u>(An et al. 2013)</u>, Norwegian fjord waters <u>(Suarez et al. 2022)</u>, river estuary sediment from China <u>(Wang et al. 2021)</u>, magnetotactic enrichment communities <u>(Lin et al. 2020)</u>, and a limonene enrichment from bog soil in Germany (Kizina et al. 2022).

### Curation

Repeat-rich regions are a common challenge for recovery of accurate and complete giant gene sequences from short read assemblies as they lead to assembly fragmentation. Thus, many genes in assemblies of the wetland soil data and public data span the entire length of their contigs. In some cases, the available protein segments have high similarity to portions of other giant proteins. In some cases, manual curation that relocates misplaced or unplaced reads and removed sequence disagreements can resolve these regions. In other cases, long read sequencing provides a means to span repeat regions. We assembled Nanopore and PacBio sequences from the same samples, in some cases producing circularized sequences. These draft genomes were curated to completion using Illumina reads to correct local errors that sometimes caused truncation of open reading frames. Curation was performed using Geneious prime 2022.1.1 (https://www.geneious.com). Alignments for illumina reads were generated using bbmap.sh using parameters specified previously in this manuscript. Alignments for pacbio and nanopore reads were generated using minimap2 <u>(Li 2018)</u>.

### Phylogeny construction

Ribosomal phylogeny in Fig. 1 was constructed using iQ-TREE v1.6.12 <u>(Hoang et al. 2018; Nguyen et al. 2015)</u> using the following parameters: -bb 1000 -m LG+G4. Sequences were isolated using GOOSOS.py (https://github.com/jwestrob/GOOSOS) utilizing PFAM models <u>(Mistry et al. 2021)</u> representing 16 syntenic ribosomal proteins <u>(Hug et al. 2016)</u>. Phylogenetic trees were visualized using iTOL <u>(Letunic and Bork 2021)</u>. *Omnitrophota* genomes were obtained from proprietary data sources as discussed above. Publicly available genomes belonging to the phyla *Planctomycetota*, *Verrucomicrobiota*, *Ratteibacteria* and *Elusimicrobiota* were obtained via BV-BRC as discussed above. *Ratteibacteria* genomes were identified through phylogeny, as they were annotated in BV-BRC as *Omnitrophica* rather than *Ratteibacteria*, although the GTDB phylum-level designations for these genomes display as *Ratteibacteria*. Treefiles are attached as **Supplementary Data S5**.

### Protein Structural Prediction

Protein structures were predicted with ColabFold <u>(Mirdita et al. 2022)</u> on a subset of the *Omnitrophota* giant proteins. Structures were visualized and aligned with ChimeraX <u>(Meng et al. 2023)</u>. Reference structures were obtained from the Protein Data Bank <u>(Berman et al. 2000)</u>.

In order to improve prediction quality, each giant protein was split into 1000 amino acid fragments and folded individually. In order to investigate potential larger-scale structure spanning multiple fragments, a 500 amino acid overlap between subsequent fragments was added. This provides both the ability to analyze structures that span more than 1,000 amino acids by aligning along the 500 aa overlap region, but also reduces the likelihood of truncating potentially co-folding regions by splitting into subfragments. Folded fragments with high average pLDDT were searched for homology to known structures using Foldseek <u>(van Kempen et al. 2023)</u>. For Dockerin-type sequences, this procedure was performed across the entire length of the giant protein sequence; for Archaeovorin sequences which contain numerous predicted transmembrane domains, regions lacking predicted transmembrane helices were folded independently. Sequential subfragments with high average pLDDT were aligned based on the overlap of amino acids in adjacent subfragments.

Protein structure files, including alignments to PDB references where applicable, are included in **Supplementary Data S6**.

### Proteomics Analysis

Decontaminated MS/MS proteomics spectra provided alongside Kizina et al. 2022 were called *de novo* using Casanovo <u>(Yilmaz et al. 2022)</u>. Alignments of called peptide fragments to the giant protein located on the *Vampirococcus* sp. LiM genome were computed using blastp <u>(Camacho et al. 2009)</u>.

### Custom Plots and Code

Heatmaps, barcharts, and scatter plots were generated using Seaborn and Matplotlib in Python. Genome diagrams (supplementary figures Sn-Sm) were generated using custom javascript and python code (**Supplementary Data S6. Custom scripts**). Custom code used to generate these figures, as well as code used in other parts of the analysis, was generated with the assistance of GPT-4.

## Data Availability

All supplementary data, including genomes used in this analysis, are available on Figshare at https://figshare.com/projects/Giant_Proteins/186507. Sequences of giant proteins, obtained from genomes in **Supplementary Data S1**, are included separately in .faa format as **Supplementary Data S8**. Newly presented genomes will be submitted to NCBI GenBank upon publication.

## Supplementary Figures

**Supplementary Figure S1.**
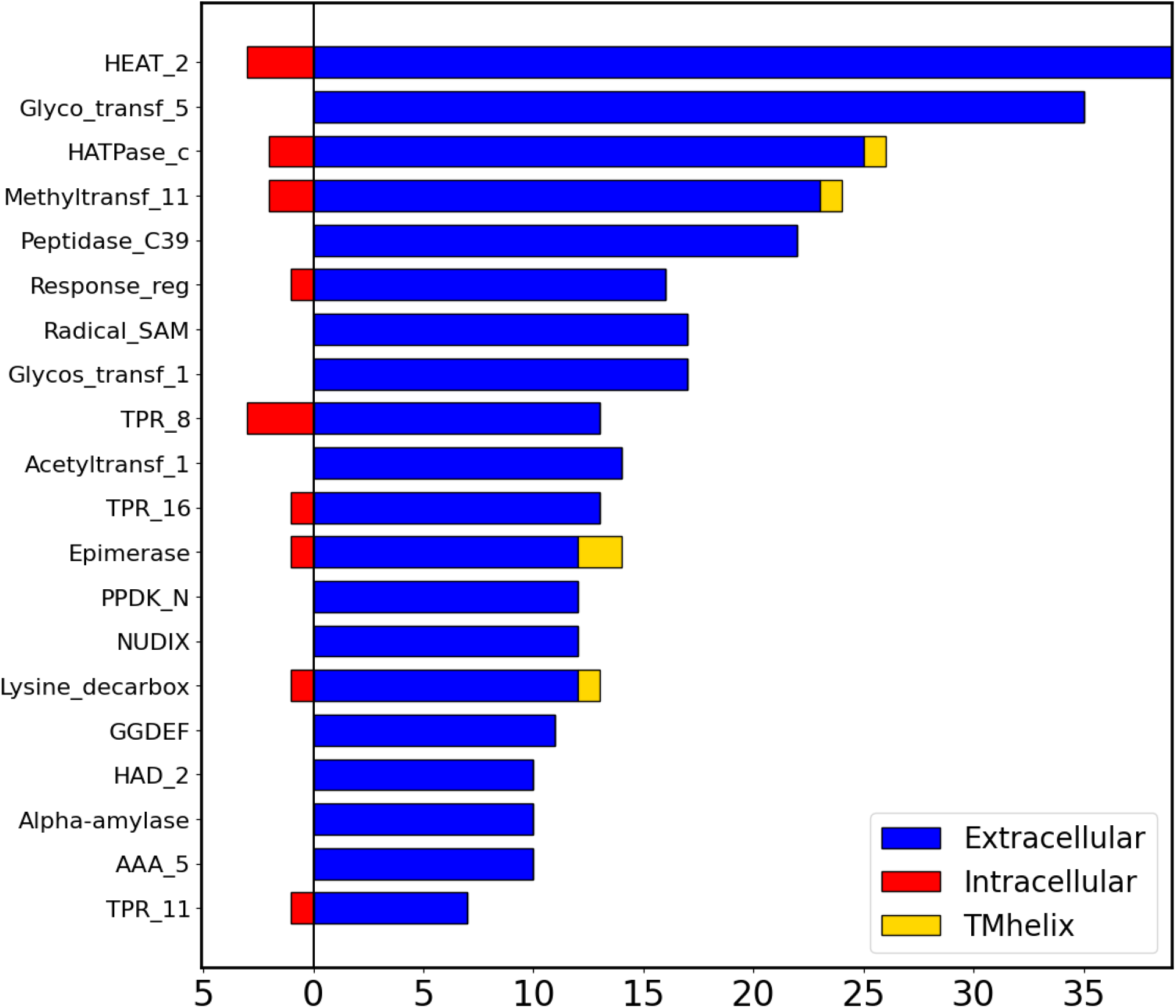
Counts of domains predicted to be exclusively extracellular (blue, right), exclusively intracellular (red, left), and domains containing transmembrane helices (gold, right) among top 20 most frequently observed domains. Localization prediction is based on the output results of TMHMM 2.0. Counts exclude those found on the three Dockerin-type giant proteins. Domain abbreviations: HEAT_2: HEAT repeats, type 2; HATPase_c: histidine kinase-like ATPase domain; Radical_SAM: SAM-dependent methyltransferase; TPR_8, TPR_11, TPR_16: Tetratricopeptide repeat, types 8, 11 and 16; PPDK_N: Pyruvate/phosphate dikinase, N-terminal domain; NUDIX: NUDIX-type hydrolase; Lysine_decarbox: putative lysine decarboxylase; GGDEF: Diguanylate cyclase; HAD_2: Haloacid dehalogenase, type 2; AAA_5: P-loop AAA-type NTPase domain, type 5.

**Supplementary Fig. S2.**
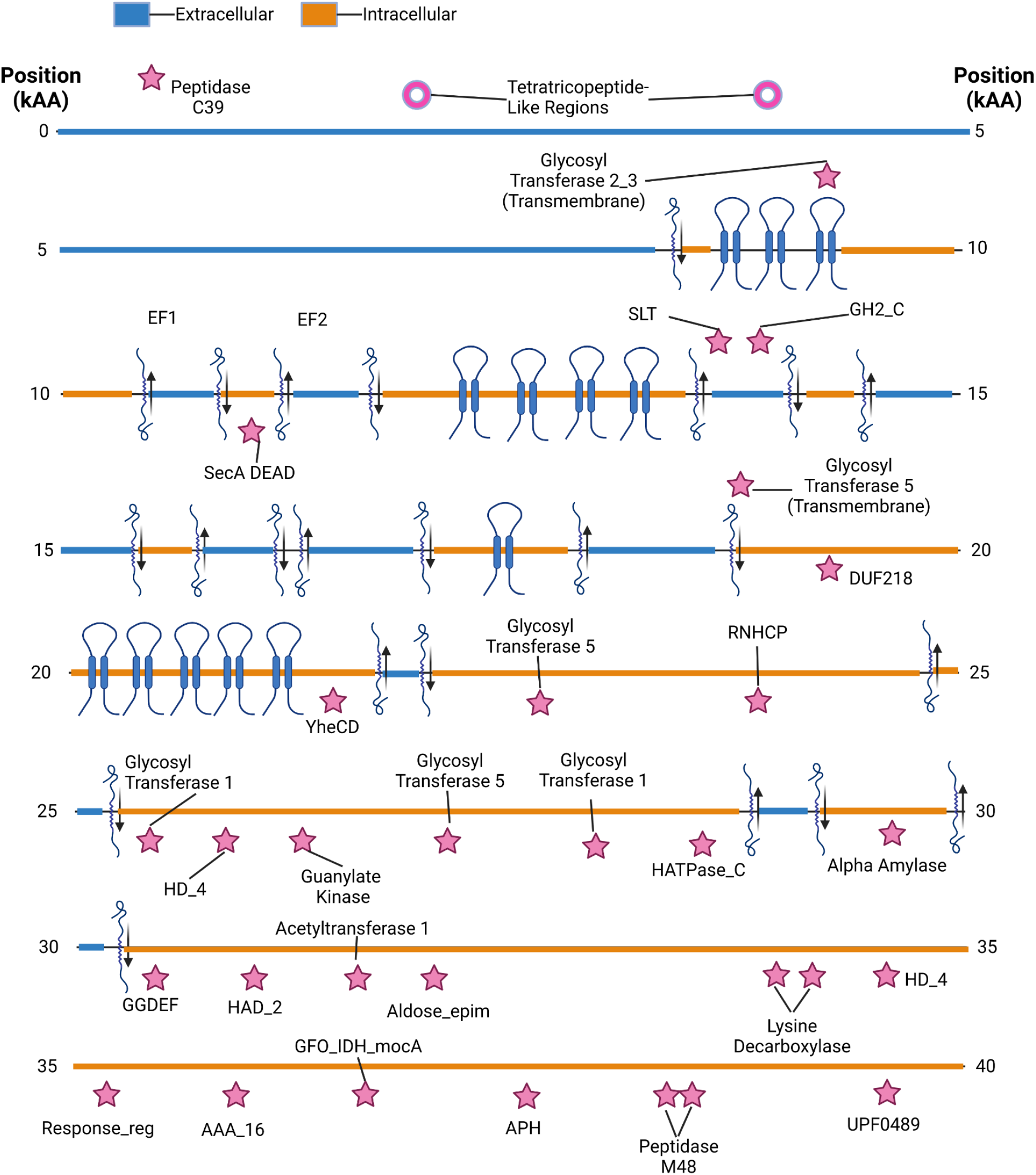
Domain diagram for giant protein found on *Velaminicoccus archaeovorus sp. LiM*. Blue and orange color indicate predicted subcellular localization. Pink stars indicate domains identified through HMM-based approaches; pink donuts indicate structures identified only through *in silico* structural analysis.

**Supplementary Fig. S3.**
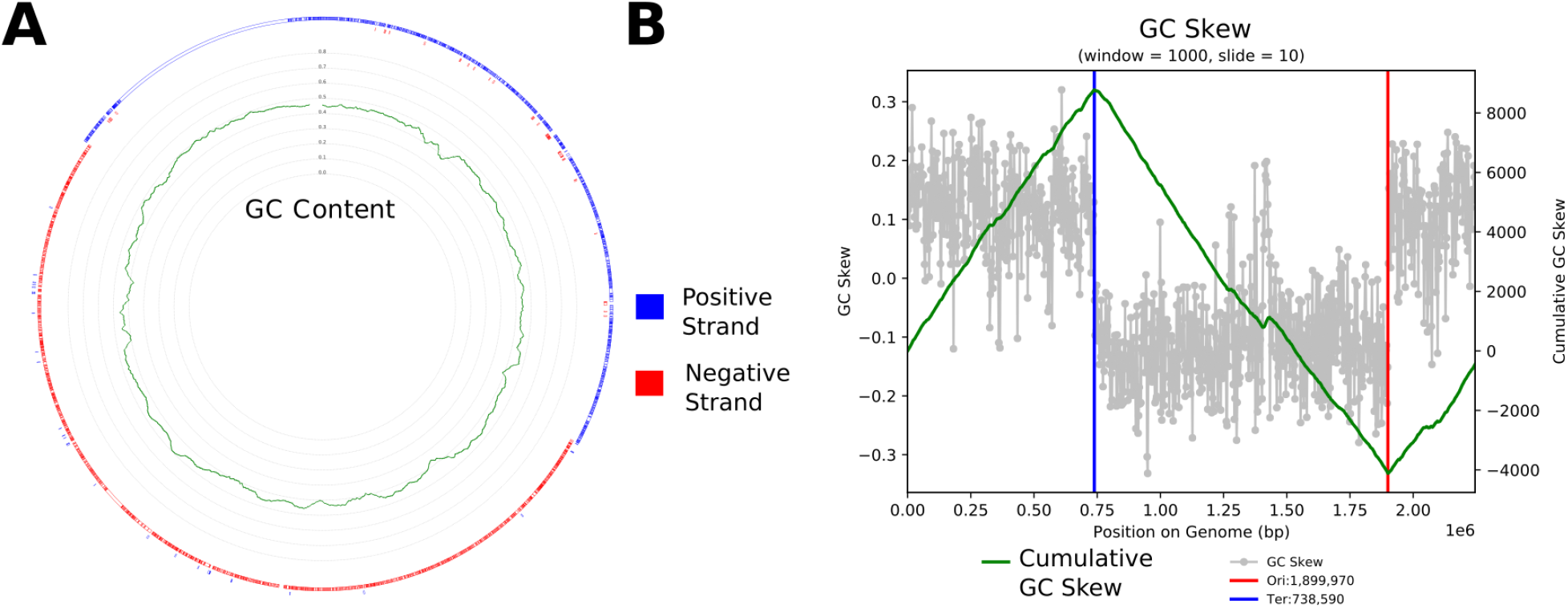
**A.** Genome diagram for curated genome SR-VP_9_9_2021_34_2B_1_4m_PACBIO-HIFI_HIFIASM-META_416_C. Diagram displays orientation of predicted open reading frames (outer rings) and GC content calculated with a four base pair moving average (interior). **B**. GC skew plot for this genome indicating the predicted origin and terminus.

**Supplementary Fig. S4.**
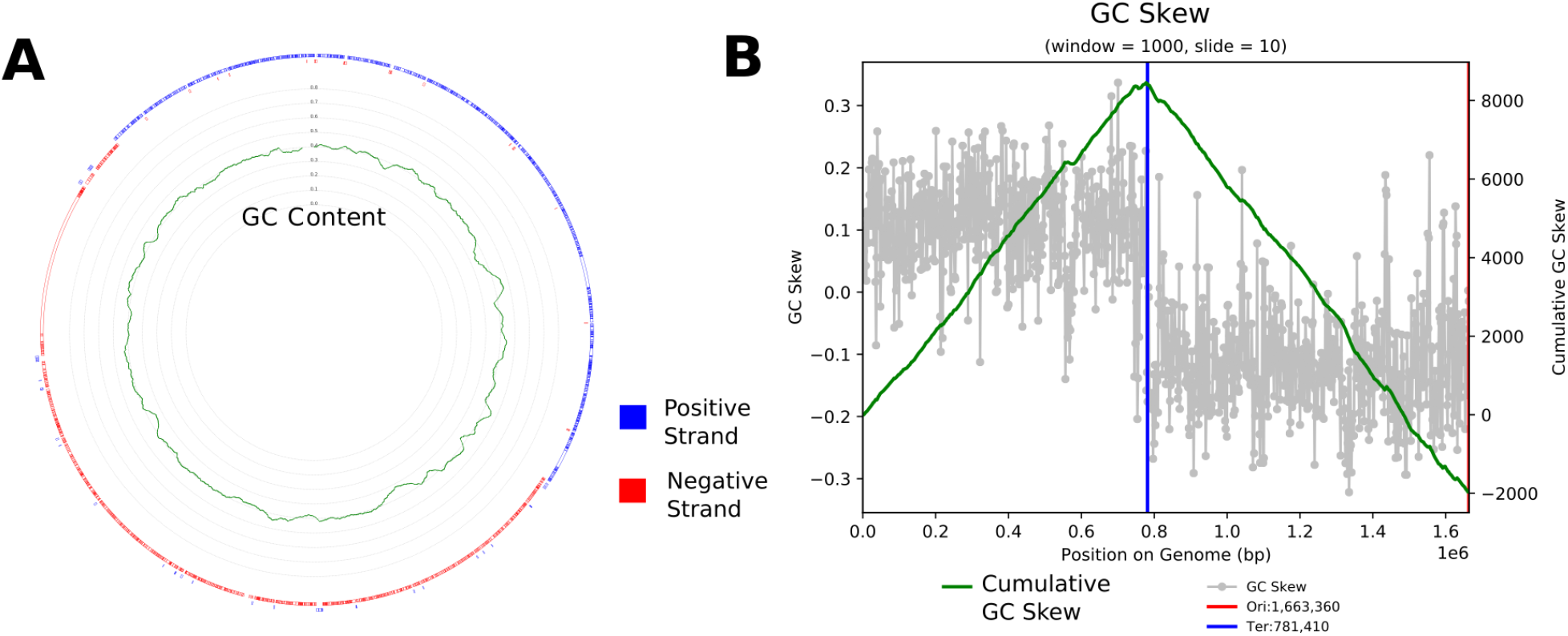
**A.** Genome diagram for curated genome SR-VP_9_9_2021_34_2B_1_4m_PACBIO-HIFI_HIFIASM-META_2055_421. Diagram displays orientation of predicted open reading frames (outer rings) and GC content calculated with a four base pair moving average (interior). **B**. GC skew plot for this genome indicating the predicted origin and terminus.

**Supplementary Fig. S5.**
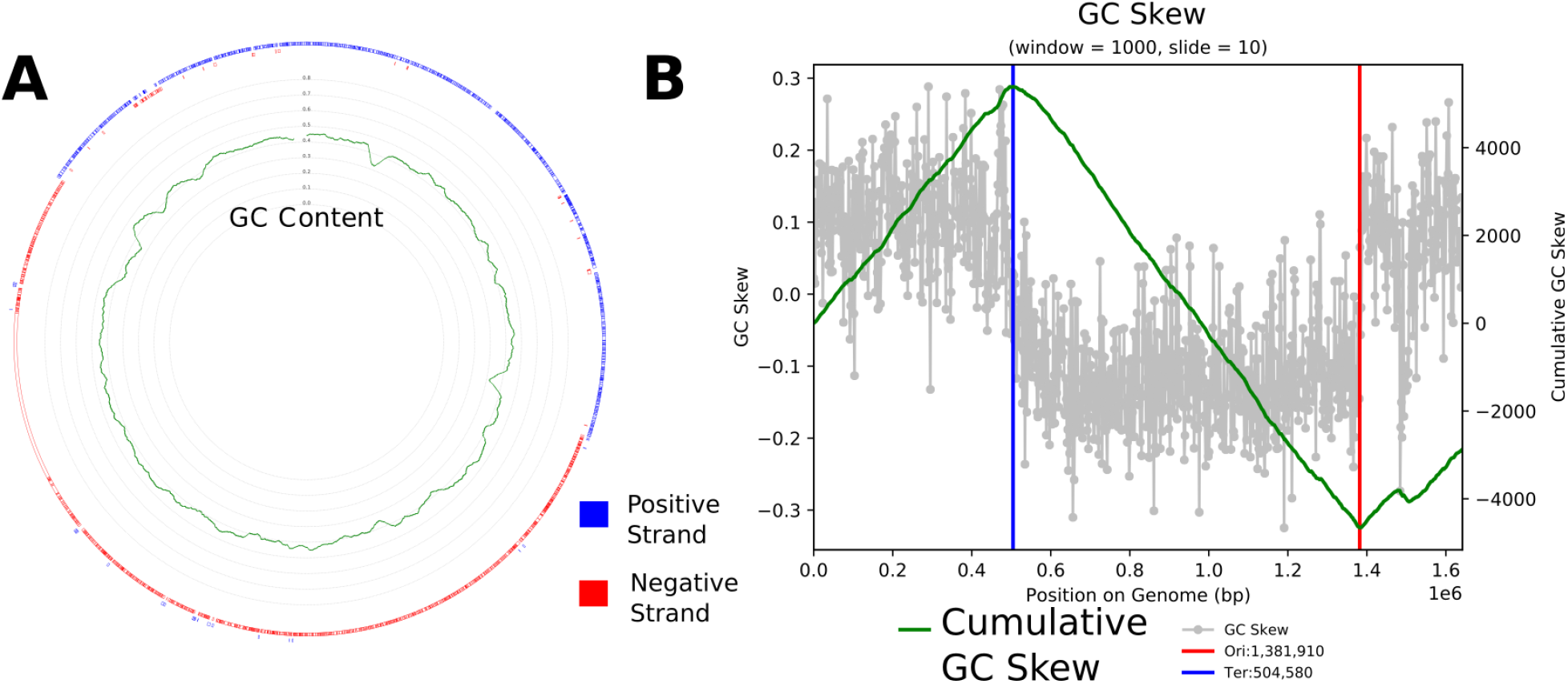
**A.** Genome diagram for curated genome SR-VP_9_9_2021_34_2B_1_4m_PACBIO-HIFI_HIFIASM-META_2055_C. Diagram displays orientation of predicted open reading frames (outer rings) and GC content calculated with a four base pair moving average (interior). **B**. GC skew plot for this genome indicating the predicted origin and terminus.

## Notes

https://figshare.com/projects/Giant_Proteins/186507

### Competing Interest Statement

The authors have declared no competing interest.

## Citations

An, Dongshan, Sean M. Caffrey, Jung Soh, Akhil Agrawal, Damon Brown, Karen Budwill, Xiaoli Dong, et al. 2013. “Metagenomics of Hydrocarbon Resource Environments Indicates Aerobic Taxa and Genes to Be Unexpectedly Common.” Environmental Science & Technology 47 (18): 10708–17.

Bell, Emma, Tiina Lamminmäki, Johannes Alneberg, Chen Qian, Weili Xiong, Robert L. Hettich, Manon Frutschi, and Rizlan Bernier-Latmani. 2022. “Active Anaerobic Methane Oxidation and Sulfur Disproportionation in the Deep Terrestrial Subsurface.” The ISME Journal 16 (6): 1583–93.

Berman, H. M., J. Westbrook, Z. Feng, G. Gilliland, T. N. Bhat, H. Weissig, I. N. Shindyalov, and P. E. Bourne. 2000. “The Protein Data Bank.” Nucleic Acids Research 28 (1): 235–42.

Bor, Batbileg, Jeffrey S. McLean, Kevin R. Foster, Lujia Cen, Thao T. To, Alejandro Serrato-Guillen, Floyd E. Dewhirst, Wenyuan Shi, and Xuesong He. 2018. “Rapid Evolution of Decreased Host Susceptibility Drives a Stable Relationship between Ultrasmall Parasite TM7x and Its Bacterial Host.” Proceedings of the National Academy of Sciences of the United States of America 115 (48): 12277–82.

Bratanis, Eleni, and Rolf Lood. 2019. “A Novel Broad-Spectrum Elastase-Like Serine Protease From the Predatory Bacterium Bdellovibrio Bacteriovorus Facilitates Elucidation of Site-Specific IgA Glycosylation Pattern.” Frontiers in Microbiology 10 (May): 971.

Buck, Moritz, Sarahi L. Garcia, Leyden Fernandez, Gaëtan Martin, Gustavo A. Martinez-Rodriguez, Jatta Saarenheimo, Jakob Zopfi, Stefan Bertilsson, and Sari Peura. 2021. “Comprehensive Dataset of Shotgun Metagenomes from Oxygen Stratified Freshwater Lakes and Ponds.” Scientific Data 8 (1): 131.

Cabello-Yeves, Pedro J., David J. Scanlan, Cristiana Callieri, Antonio Picazo, Lena Schallenberg, Paula Huber, Juan J. Roda-Garcia, et al. 2022. “α-Cyanobacteria Possessing Form IA RuBisCO Globally Dominate Aquatic Habitats.” The ISME Journal 16 (10): 2421–32.

Camacho, Christiam, George Coulouris, Vahram Avagyan, Ning Ma, Jason Papadopoulos, Kevin Bealer, and Thomas L. Madden. 2009. “BLAST+: Architecture and Applications.” BMC Bioinformatics 10 (December): 421.

Carvalho, Ana L., Fernando M. V. Dias, José A. M. Prates, Tibor Nagy, Harry J. Gilbert, Gideon J. Davies, Luís M. A. Ferreira, Maria J. Romão, and Carlos M. G. A. Fontes. 2003. “Cellulosome Assembly Revealed by the Crystal Structure of the Cohesin-Dockerin Complex.” Proceedings of the National Academy of Sciences of the United States of America 100 (24): 13809–14.

Casar, C. P., L. M. Momper, B. R. Kruger, and M. R. Osburn. 2021. “Iron-Fueled Life in the Continental Subsurface: Deep Mine Microbial Observatory, South Dakota, USA.” Applied and Environmental Microbiology 87 (20): e0083221.

Cerveny, Lukas, Adela Straskova, Vera Dankova, Anetta Hartlova, Martina Ceckova, Frantisek Staud, and Jiri Stulik. 2013. “Tetratricopeptide Repeat Motifs in the World of Bacterial Pathogens: Role in Virulence Mechanisms.” Infection and Immunity 81 (3): 629–35.

Chaussonnerie, Sébastien, Pierre-Loïc Saaidi, Edgardo Ugarte, Agnès Barbance, Aurélie Fossey, Valérie Barbe, Gabor Gyapay, et al. 2016. “Microbial Degradation of a Recalcitrant Pesticide: Chlordecone.” Frontiers in Microbiology 7 (December): 2025.

Dai, Shuyan, Cancan Sun, Kemin Tan, Sheng Ye, and Rongguang Zhang. 2017. “Structure of Thrombospondin Type 3 Repeats in Bacterial Outer Membrane Protein A Reveals Its Intra-Repeat Disulfide Bond-Dependent Calcium-Binding Capability.” Cell Calcium 66 (September): 78–89.

Davidov, Yaacov, Dorothee Huchon, Susan F. Koval, and Edouard Jurkevitch. 2006. “A New Alpha-Proteobacterial Clade of Bdellovibrio-like Predators: Implications for the Mitochondrial Endosymbiotic Theory.” Environmental Microbiology 8 (12): 2179–88.

Elcheninov, Alexander G., Peter Menzel, Soley R. Gudbergsdottir, Alexei I. Slesarev, Vitaly V. Kadnikov, Anders Krogh, Elizaveta A. Bonch-Osmolovskaya, Xu Peng, and Ilya V. Kublanov. 2017. “Sugar Metabolism of the First Thermophilic Planctomycete Thermogutta Terrifontis: Comparative Genomic and Transcriptomic Approaches.” Frontiers in Microbiology 8 (November): 2140.

Feng, Xiaowen, Haoyu Cheng, Daniel Portik, and Heng Li. 2022. “Metagenome Assembly of High-Fidelity Long Reads with Hifiasm-Meta.” Nature Methods 19 (6): 671–74.

Flaugnatti, Nicolas, Chiara Rapisarda, Martial Rey, Solène G. Beauvois, Viet Anh Nguyen, Stéphane Canaan, Eric Durand, et al. 2020. “Structural Basis for Loading and Inhibition of a Bacterial T6SS Phospholipase Effector by the VgrG Spike.” The EMBO Journal 39 (11): e104129.

Frostl, J. M., and J. Overmann. 1998. “Physiology and Tactic Response of the Phototrophic Consortium ‘Chlorochromatium Aggregatum.’” Archives of Microbiology 169 (2): 129–35.

Grégoire, Daniel S., Nikhil A. George, and Laura A. Hug. 2023. “Microbial Methane Cycling in a Landfill on a Decadal Time Scale.” bioRxiv. 10.1101/2023.01.20.524919.

Gupta, Shilpi, Chi Tang, Michael Tran, and Daniel E. Kadouri. 2016. “Effect of Predatory Bacteria on Human Cell Lines.” PloS One 11 (8): e0161242.

Haas, W., and J. A. Banas. 2000. “Ligand-Binding Properties of the Carboxyl-Terminal Repeat Domain of Streptococcus Mutans Glucan-Binding Protein A.” Journal of Bacteriology 182 (3): 728–33.

Hallgren, Jeppe, Konstantinos D. Tsirigos, Mads Damgaard Pedersen, José Juan Almagro Armenteros, Paolo Marcatili, Henrik Nielsen, Anders Krogh, and Ole Winther. 2022. “DeepTMHMM Predicts Alpha and Beta Transmembrane Proteins Using Deep Neural Networks.” bioRxiv. 10.1101/2022.04.08.487609.

Hoang, Diep Thi, Olga Chernomor, Arndt von Haeseler, Bui Quang Minh, and Le Sy Vinh. 2018. “UFBoot2: Improving the Ultrafast Bootstrap Approximation.” Molecular Biology and Evolution 35 (2): 518–22.

Hug, Laura A., Brett J. Baker, Karthik Anantharaman, Christopher T. Brown, Alexander J. Probst, Cindy J. Castelle, Cristina N. Butterfield, et al. 2016. “A New View of the Tree of Life.” Nature Microbiology 1 (April): 16048.

Hunt, John F., Sevil Weinkauf, Lisa Henry, John J. Fak, Paul McNicholas, Donald B. Oliver, and Johann Deisenhofer. 2002. “Nucleotide Control of Interdomain Interactions in the Conformational Reaction Cycle of SecA.” Science 297 (5589): 2018–26.

Hurst, Mark R. H., Sandra A. Jones, Tan Binglin, Lincoln A. Harper, Trevor A. Jackson, and Travis R. Glare. 2011. “The Main Virulence Determinant of Yersinia Entomophaga MH96 Is a Broad-Host-Range Toxin Complex Active against Insects.” Journal of Bacteriology 193 (8): 1966–80.

Hyatt, Doug, Gwo-Liang Chen, Philip F. Locascio, Miriam L. Land, Frank W. Larimer, and Loren J. Hauser. 2010. “Prodigal: Prokaryotic Gene Recognition and Translation Initiation Site Identification.” BMC Bioinformatics 11 (March): 119.

Ikemura, H., and M. Inouye. 1988. “In Vitro Processing of pro-Subtilisin Produced in Escherichia Coli.” The Journal of Biological Chemistry 263 (26): 12959–63.

Iyer, Sai Prasad N., and Gerald W. Hart. 2003. “Roles of the Tetratricopeptide Repeat Domain in O-GlcNAc Transferase Targeting and Protein Substrate Specificity.” The Journal of Biological Chemistry 278 (27): 24608–16.

Kang, Dongwan D., Feng Li, Edward Kirton, Ashleigh Thomas, Rob Egan, Hong An, and Zhong Wang. 2019. “MetaBAT 2: An Adaptive Binning Algorithm for Robust and Efficient Genome Reconstruction from Metagenome Assemblies.” PeerJ 7 (July): e7359.

Kempen, Michel van, Stephanie S. Kim, Charlotte Tumescheit, Milot Mirdita, Jeongjae Lee, Cameron L. M. Gilchrist, Johannes Söding, and Martin Steinegger. 2023. “Fast and Accurate Protein Structure Search with Foldseek.” *Nature Biotechnology*, May. 10.1038/s41587-023-01773-0.

Kizina Jana, Jordan Sebastian F. A., Martens Gerrit Alexander, Lonsing Almud, Probian Christina, Kolovou Androniki, Santarella-Mellwig Rachel, et al. 2022. “Methanosaeta and ‘Candidatus Velamenicoccus Archaeovorus.’” Applied and Environmental Microbiology 88 (7): e02407–21.

Krogh, Anders, Björn Larsson, Gunnar von Heijne, and Erik L. L. Sonnhammer. 2001. “Predicting Transmembrane Protein Topology with a Hidden Markov Model: Application to Complete genomes11Edited by F. Cohen.” Journal of Molecular Biology 305 (3): 567–80.

Kumbhar, Charushila, Praneitha Mudliar, Latika Bhatia, Aseem Kshirsagar, and Milind Watve. 2014. “Widespread Predatory Abilities in the Genus Streptomyces.” Archives of Microbiology 196 (4): 235–48.

Kuwabara, Naoyuki, Hiroshi Manya, Takeyuki Yamada, Hiroaki Tateno, Motoi Kanagawa, Kazuhiro Kobayashi, Keiko Akasaka-Manya, et al. 2016. “Carbohydrate-Binding Domain of the POMGnT1 Stem Region Modulates O-Mannosylation Sites of α-Dystroglycan.” Proceedings of the National Academy of Sciences of the United States of America 113 (33): 9280–85.

Kvansakul, Marc, Josephine C. Adams, and Erhard Hohenester. 2004. “Structure of a Thrombospondin C-Terminal Fragment Reveals a Novel Calcium Core in the Type 3 Repeats.” The EMBO Journal 23 (6): 1223–33.

Lecher, Justin, Christian K. W. Schwarz, Matthias Stoldt, Sander H. J. Smits, Dieter Willbold, and Lutz Schmitt. 2012. “An RTX Transporter Tethers Its Unfolded Substrate during Secretion via a Unique N-Terminal Domain.” Structure 20 (10): 1778–87.

Letunic, Ivica, and Peer Bork. 2021. “Interactive Tree Of Life (iTOL) v5: An Online Tool for Phylogenetic Tree Display and Annotation.” Nucleic Acids Research 49 (W1): W293–96.

Li, Dinghua, Chi-Man Liu, Ruibang Luo, Kunihiko Sadakane, and Tak-Wah Lam. 2015. “MEGAHIT: An Ultra-Fast Single-Node Solution for Large and Complex Metagenomics Assembly via Succinct de Bruijn Graph.” Bioinformatics 31 (10): 1674–76.

Lin, Wei, Wensi Zhang, Greig A. Paterson, Qiyun Zhu, Xiang Zhao, Rob Knight, Dennis A. Bazylinski, Andrew P. Roberts, and Yongxin Pan. 2020. “Expanding Magnetic Organelle Biogenesis in the Domain Bacteria.” Microbiome 8 (1): 152.

Liu, Lei, Yulin Wang, Yu Yang, Depeng Wang, Suk Hang Cheng, Chunmiao Zheng, and Tong Zhang. 2021. “Charting the Complexity of the Activated Sludge Microbiome through a Hybrid Sequencing Strategy.” Microbiome 9 (1): 205.

Liu, Zhenfeng, Johannes Müller, Tao Li, Richard M. Alvey, Kajetan Vogl, Niels-Ulrik Frigaard, Nathan C. Rockwell, et al. 2013. “Genomic Analysis Reveals Key Aspects of Prokaryotic Symbiosis in the Phototrophic Consortium ‘Chlorochromatium Aggregatum.’” Genome Biology 14 (11): R127.

Livingstone, Paul G., Russell M. Morphew, Alan R. Cookson, and David E. Whitworth. 2018. “Genome Analysis, Metabolic Potential, and Predatory Capabilities of Herpetosiphon Llansteffanense Sp. Nov.” Applied and Environmental Microbiology 84 (22). 10.1128/AEM.01040-18.

Mathieu, Sophie V., Karoline Saboia Aragão, Anne Imberty, and Annabelle Varrot. 2010. “Discoidin I from Dictyostelium Discoideum and Interactions with Oligosaccharides: Specificity, Affinity, Crystal Structures, and Comparison with Discoidin II.” Journal of Molecular Biology 400 (3): 540–54.

Mei, Ran, and Wen-Tso Liu. 2022. “Meta-Omics-Supervised Characterization of Respiration Activities Associated with Microbial Immigrants in Anaerobic Sludge Digesters.” Environmental Science & Technology 56 (10): 6689–98.

Meng, Elaine C., Thomas D. Goddard, Eric F. Pettersen, Greg S. Couch, Zach J. Pearson, John H. Morris, and Thomas E. Ferrin. 2023. “UCSF ChimeraX: Tools for Structure Building and Analysis.” Protein Science: A Publication of the Protein Society 32 (11): e4792.

Meshcheryakov, Vladimir A., Satoshi Shibata, Makoto Tokoro Schreiber, Alejandro Villar-Briones, Kenneth F. Jarrell, Shin-Ichi Aizawa, and Matthias Wolf. 2019. “High-Resolution Archaellum Structure Reveals a Conserved Metal-Binding Site.” EMBO Reports 20 (5). 10.15252/embr.201846340.

Mirdita, Milot, Konstantin Schütze, Yoshitaka Moriwaki, Lim Heo, Sergey Ovchinnikov, and Martin Steinegger. 2022. “ColabFold: Making Protein Folding Accessible to All.” Nature Methods 19 (6): 679–82.

Mistry, Jaina, Sara Chuguransky, Lowri Williams, Matloob Qureshi, Gustavo A. Salazar, Erik L. L. Sonnhammer, Silvio C. E. Tosatto, et al. 2021. “Pfam: The Protein Families Database in 2021.” Nucleic Acids Research 49 (D1): D412–19.

Mitchell, Alex L., Maxim Scheremetjew, Hubert Denise, Simon Potter, Aleksandra Tarkowska, Matloob Qureshi, Gustavo A. Salazar, et al. 2018. “EBI Metagenomics in 2017: Enriching the Analysis of Microbial Communities, from Sequence Reads to Assemblies.” Nucleic Acids Research 46 (D1): D726–35.

Montanier, Cedric, Alicia Lammerts van Bueren, Claire Dumon, James E. Flint, Marcia A. Correia, Jose A. Prates, Susan J. Firbank, et al. 2009. “Evidence That Family 35 Carbohydrate Binding Modules Display Conserved Specificity but Divergent Function.” Proceedings of the National Academy of Sciences of the United States of America 106 (9): 3065–70.

Moreira, David, Yvan Zivanovic, Ana I. López-Archilla, Miguel Iniesto, and Purificación López-García. 2021. “Reductive Evolution and Unique Predatory Mode in the CPR Bacterium Vampirococcus Lugosii.” Nature Communications 12 (1): 2454.

Mosley, Olivia E., Emilie Gios, Louise Weaver, Murray Close, Chris Daughney, Rob van der Raaij, Heather Martindale, and Kim M. Handley. 2022. “Metabolic Diversity and Aero-Tolerance in Anammox Bacteria from Geochemically Distinct Aquifers.” mSystems 7 (1): e0125521.

Neuwald, A. F., and T. Hirano. 2000. “HEAT Repeats Associated with Condensins, Cohesins, and Other Complexes Involved in Chromosome-Related Functions.” Genome Research 10 (10): 1445–52.

Nguyen, Lam-Tung, Heiko A. Schmidt, Arndt von Haeseler, and Bui Quang Minh. 2015. “IQ-TREE: A Fast and Effective Stochastic Algorithm for Estimating Maximum-Likelihood Phylogenies.” Molecular Biology and Evolution 32 (1): 268–74.

Nissen, Jakob Nybo, Joachim Johansen, Rosa Lundbye Allesøe, Casper Kaae Sønderby, Jose Juan Almagro Armenteros, Christopher Heje Grønbech, Lars Juhl Jensen, et al. 2021. “Improved Metagenome Binning and Assembly Using Deep Variational Autoencoders.” Nature Biotechnology 39 (5): 555–60.

Pagès, S., A. Bélaïch, J. P. Bélaïch, E. Morag, R. Lamed, Y. Shoham, and E. A. Bayer. 1997. “Species-Specificity of the Cohesin-Dockerin Interaction between Clostridium Thermocellum and Clostridium Cellulolyticum: Prediction of Specificity Determinants of the Dockerin Domain.” Proteins 29 (4): 517–27.

Peng, Yu, Henry C. M. Leung, S. M. Yiu, and Francis Y. L. Chin. 2012. “IDBA-UD: A de Novo Assembler for Single-Cell and Metagenomic Sequencing Data with Highly Uneven Depth.” Bioinformatics 28 (11): 1420–28.

Pereira, Joana, and Andrei N. Lupas. 2022. “New β-Propellers Are Continuously Amplified From Single Blades in All Major Lineages of the β-Propeller Superfamily.” Frontiers in Molecular Biosciences 9 (June): 895496.

Perez-Molphe-Montoya, Eugenio, Kirsten Küsel, and Will A. Overholt. 2022. “Redefining the Phylogenetic and Metabolic Diversity of Phylum Omnitrophota.” Environmental Microbiology 24 (11): 5437–49.

Plaza Onate, Florian, Camille Coulibaly, Caroline Achard, Amine Ghozlane, Nicolas Pons, Etienne Ruppe, Benoit Dile, et al. 2022. “MicroReset: Characterization of the Rabbit (Oryctolagus Cuniculus) Fecal Metagenome and Resistome by Deep Shotgun Sequencing.” Recherche Data Gouv. 10.15454/5EJKAS.

Rashidan, K. K., and D. F. Bird. 2001. “Role of Predatory Bacteria in the Termination of a Cyanobacterial Bloom.” Microbial Ecology 41 (2): 97–105.

Rawlings, Neil D., Alan J. Barrett, Paul D. Thomas, Xiaosong Huang, Alex Bateman, and Robert D. Finn. 2018. “The MEROPS Database of Proteolytic Enzymes, Their Substrates and Inhibitors in 2017 and a Comparison with Peptidases in the PANTHER Database.” Nucleic Acids Research 46 (D1): D624–32.

Reva, Oleg, and Burkhard Tümmler. 2008. “Think Big--Giant Genes in Bacteria.” Environmental Microbiology 10 (3): 768–77.

Rincon, Marco T., Shi-You Ding, Sheila I. McCrae, Jennifer C. Martin, Vincenzo Aurilia, Raphael Lamed, Yuval Shoham, Edward A. Bayer, and Harry J. Flint. 2003. “Novel Organization and Divergent Dockerin Specificities in the Cellulosome System of Ruminococcus Flavefaciens.” Journal of Bacteriology 185 (3): 703–13.

Schoelmerich, Marie C., Lynn Ly, Jacob West-Roberts, Ling-Dong Shi, Cong Shen, Nikhil S. Malvankar, Najwa Taib, et al. 2023. “Borg Extrachromosomal Elements of Methane-Oxidizing Archaea Have Conserved and Expressed Genetic Repertoires.” bioRxiv. 10.1101/2023.08.01.549754.

Seymour, Cale O., Marike Palmer, Eric D. Becraft, Ramunas Stepanauskas, Ariel D. Friel, Frederik Schulz, Tanja Woyke, et al. 2023. “Hyperactive Nanobacteria with Host-Dependent Traits Pervade Omnitrophota.” Nature Microbiology 8 (4): 727–44.

Shiratori, Takashi, Shigekatsu Suzuki, Yukako Kakizawa, and Ken-Ichiro Ishida. 2019. “Phagocytosis-like Cell Engulfment by a Planctomycete Bacterium.” Nature Communications 10 (1): 5529.

Sieber, Christian M. K., Alexander J. Probst, Allison Sharrar, Brian C. Thomas, Matthias Hess, Susannah G. Tringe, and Jillian F. Banfield. 2018. “Recovery of Genomes from Metagenomes via a Dereplication, Aggregation and Scoring Strategy.” Nature Microbiology 3 (7): 836–43.

Stevens, Tim J., and Max Paoli. 2008. “RCC1-like Repeat Proteins: A Pangenomic, Structurally Diverse New Superfamily of Beta-Propeller Domains.” Proteins 70 (2): 378–87.

Suarez, Carolina, Paula Dalcin Martins, Mike S. M. Jetten, Sabina Karačić, Britt Marie Wilén, Oskar Modin, Per Hagelia, Malte Hermansson, and Frank Persson. 2022. “Metagenomic Evidence of a Novel Family of Anammox Bacteria in a Subsea Environment.” Environmental Microbiology 24 (5): 2348–60.

Swart, Anna Leoni, Laura Gomez-Valero, Carmen Buchrieser, and Hubert Hilbi. 2020. “Evolution and Function of Bacterial RCC1 Repeat Effectors.” Cellular Microbiology 22 (10): e13246.

Thiery, Susanne, and Christine Kaimer. 2020. “The Predation Strategy of Myxococcus Xanthus.” Frontiers in Microbiology 11 (January): 2.

UniProt Consortium. 2023. “UniProt: The Universal Protein Knowledgebase in 2023.” Nucleic Acids Research 51 (D1): D523–31.

Van Essche, M., M. Quirynen, I. Sliepen, G. Loozen, N. Boon, J. Van Eldere, and W. Teughels. 2011. “Killing of Anaerobic Pathogens by Predatory Bacteria.” Molecular Oral Microbiology 26 (1): 52–61.

Wang, Wenxiu, Jianchang Tao, Ke Yu, Chen He, Jianjun Wang, Penghui Li, Hongmei Chen, Bu Xu, Quan Shi, and Chuanlun Zhang. 2021. “Vertical Stratification of Dissolved Organic Matter Linked to Distinct Microbial Communities in Subtropic Estuarine Sediments.” Frontiers in Microbiology 12 (July): 697860.

Wu, Yu-Wei, Blake A. Simmons, and Steven W. Singer. 2016. “MaxBin 2.0: An Automated Binning Algorithm to Recover Genomes from Multiple Metagenomic Datasets.” Bioinformatics 32 (4): 605–7.

Yilmaz, Melih, William Fondrie, Wout Bittremieux, Sewoong Oh, and William S. Noble. 17--23 Jul 2022. “De Novo Mass Spectrometry Peptide Sequencing with a Transformer Model.” In Proceedings of the 39th International Conference on Machine Learning, edited by Kamalika Chaudhuri, Stefanie Jegelka, Le Song, Csaba Szepesvari, Gang Niu, and Sivan Sabato, 162:25514–22. Proceedings of Machine Learning Research. PMLR.

